# Novel strategies of Raman imaging for monitoring intracellular retinoid metabolism in cancer cells

**DOI:** 10.1101/2020.05.05.078410

**Authors:** Halina Abramczyk, Anna Imiela, Jakub Surmacki

**Affiliations:** Lodz University of Technology, Faculty of Chemistry, Institute of Applied Radiation Chemistry, Laboratory of Laser Molecular Spectroscopy, Wroblewskiego 15, 93-590 Lodz, Poland

**Keywords:** Raman spectroscopy, Raman imaging, U-87 MG cell lines, astrocytes, brain cancer, glioblastoma, phosphorylation, retinoids, retinol binding protein

## Abstract

We developed a label-free Raman method for whole-cell biochemical imaging to detect molecular processes that occur in normal and cancer brain cells due to retinol transport in human cancers at the level of isolated organelles. Our approach allows to create biochemical maps of retinoids localization in lipid droplets, mitochondria and nuclei in single cells. The maps were capable of discriminating triglycerides (TAG) from retinoids (RE) in lipid droplets (LD), and mitochondria providing an excellent tool to monitor intracellular retinoid metabolism. We detected spectral changes that arose in proteins and lipids due to retinoid metabolism in human cell lines of normal astrocytes and high-grade cancer cells of glioblastoma as well as in human medulloblastoma and glioblastoma tissue. Raman imaging is an effective tool for monitoring retinoids and retinol binding proteins involved in carcinogenesis by detecting unique spectral signatures of vibrations. We found two functionally distinct lipid droplets: TAG-LD, for energy storage, and RE-LD, for regulating mechanisms of signal transduction. Raman polarization measurements revealed the occurrence of conformational changes affecting discrete regions of proteins associated with retinol binding. Aberrant expression of retinoids and retinol binding proteins in human tumours were localized in lipid droplets, and mitochondria.

## Introduction

Vitamin A plays multiple functions in a wide range of biological processes including cell differentiation, proliferation, and apoptosis. Recent results analyzed in the context of a number of literature studies suggest that processes that occur in carotenoids and retinoids play an important role in the molecular mechanisms of carcinogenesis.[1–5] In the past, considerable effort has been directed towards the cancer chemoprevention properties of carotenoids and retinoids[6,7], but relatively little attention has been paid to their role in molecular mechanisms of cancer development. Although effort has been focused almost exclusively on their antioxidant activity, their biological activity goes far beyond this signature, and a careful examination of other possible involvements in metabolism and signaling of cells is needed. It has recently been reported that carotenoid level is one of the most common abnormalities in human cancers. Carotenoid concentration is lower in human cancers than in normal cells.[8–10] In recent years, new results have been emerged indicating a role of retinoids in lipid droplets.[11,12] It has recently been reported that retinyl esters are replaced by polyunsaturated triacylglycerol species in lipid droplets of hepatic stellate cells during activation[11] and in breast and brain cancer cells.[12]

Carotenoids are delivered to the human body via the diet, because they are present in a wide range of vegetables and fruits.[13] Some carotenoids (β-carotene, α-carotene and β-cryptoxanthin) belong to the family of provitamin A carotenoids because they are precursors of vitamin A, a fat-soluble retinol.[14] Beta-carotene is transformed by enzymatically cleavage into 2 molecules of retinaldehyde, which can then can be either oxidized to the active form of vitamin A (retinoic acid), or reduced to retinal.[15]

Vitamin A is also delivered directly as preformed vitamin A, which is naturally present in food of animal origin such as meat, fish, and dairy products[16], consisting primarily of retinyl palmitate (fatty acid ester), and retinol, and in much smaller concentrations as retinoic acid. The preformed vitamin A is provided in food of animal origin via the absorption of retinol from the alimentary tract accompanied by the absorption of other food molecules. The retinol enzymatically conjugates to some of the absorbed molecules, particularly fatty acids to form esters such as retinyl palmitate, which is also converted to retinol within the lumen of the small intestine and then re-esterified to form retinyl ester within the enterocyte.[17] After the conversion of carotenoids and retinyl esters, retinol is transported through the lymphatic or blood circulation to other cells and tissues where it binds to retinol-binding proteins (RBP) in the liver, controlling the release of retinol from storage when it is needed.[18–20] A large number of physiological processes, such as vision, epithelial differentiation in cancers, inflammatory processes, and spermatogenesis, are attributed to vitamin A.[21–32]

The involvement of vitamin A in processes of vision is its most well-known characteristic, but other mechanisms linked to vitamin A are much less well understood and await further investigation. The relationship between retinoids and cancer has been reported in many studies.[1,27,33] Several studies using animal models have reported an inhibitory role of retinoids in breast cancer.[1,25] Lower expression of cellular retinol-binding protein-I (CRBP-1) has been found in breast and other human cancers, but its significance is not well understood.[1] It has been reported that retinoids can be used in the treatment of glioma.[33] Alterations in cellular binding protein expression modify retinoid biological functions and result in abnormal glucose and energy metabolism, thus increasing the predisposition to breast cancer, prostate cancer, ovarian adenocarcinoma, and glioblastoma.[34] Vitamin A has been reported to control oxidative phosphorylation in mitochondria.[35] As it is well known, the most universal feature of cancer is a high rate of glucose uptake and enhanced glycolytic activities followed by lactate fermentation known as the Warburg effect.[36] Understanding the mechanism of altered energy metabolism that replaces the more efficient oxidative phosphorylation in terms of the yield of adenosine triphosphate (ATP) with the less efficient glycolysis represents the pivotal problem that must be solved for progress in eliminating cancer. During the past decade, many studies have suggested that the switch from oxidative phosphorylation to lactate fermentation is controlled by the metabolic fate of pyruvate regulated by tyrosine phosphorylation.[37–39]

Vitamin A plays an important role in cell signaling. Abnormal retinoid signaling in the cytoplasm and nucleus has been reported in human cancers.[40] Vitamin A is involved in cell signaling by attachment to the retinol-binding protein receptor termed STRA6 (stimulated by retinoic acid 6).[41] STRA6 is a transmembrane cell surface protein that functions as a receptor for a family of retinol-binding proteins (RBPs) that play roles as carrier proteins that bind to retinol.[2,42] STRA6 is a ligand-activated cell surface signaling receptor that upon binding of the retinol to RBP (also termed the holo-RBP or RBP-ROH complex) activates JAK/STAT signaling to transfer information from the extracellular environment to the nucleus, resulting in DNA transcription and inducing the expression of target genes involved in many processes such as immunity, proliferation, differentiation, and apoptosis. Disrupted JAK/STAT signaling may lead to a variety of diseases, such as cancers and disorders of the immune system.[43] Janus kinases (JAKs) belong to an enzyme family of intracellular, nonreceptor tyrosine kinases that activate STAT signaling. STAT (signal transducer and activator of transcription) is a member of protein family that activates genes through transcription. The transport of retinol to the interior of the cell occurs via the retinol acceptor CRBP1 protein attached to the CRBP binding loop of STRA6. This event activates the intracellular non-receptor tyrosine kinase enzyme JAK2, which transfers a phosphate group from ATP to a protein in the cell, thereby phosphorylating STRA6 at Y643 (tyrosine).[2] The detailed mechanism of JAK2 activation by the retinol-RBP complex is still unknown and awaits further investigation to elucidate the cascade of downstream signals and events necessary to activate (or suppress) gene transcription in the nucleus.

To avoid the pitfalls of conventional approaches and to obtain biochemical information, it is necessary to develop new tools for the precise analysis of signals from molecules present in specific organelles of cells: proteins, fatty acids and lipids, DNA, RNA, carbohydrates, carotenoids, retinoids, and primary metabolites, which can be obtained by Raman microscopy.

Raman microspectroscopy - a spectroscopic technique based on inelastic scattering of monochromatic light - does not require labelling of the molecules of interest and enables direct specific chemical imaging of biomolecules such as DNA/RNA, proteins, and lipids in intact cells and tissues.[44–46]

We will show that label-free Raman microscopy could also enable the clarification of the precise role of retinoids in the metabolism and signaling of cancer cells. The purpose of this paper is to examine the role of retinoids in human cell lines of normal astrocytes (NHA) and high-grade tumor cells of glioblastoma (U-87 MG) and in human medulloblastoma and glioblastoma tissues.

This paper provides a basis for substantial revision of the previous interpretation of the biochemical events associated with retinol in normal and cancer cells. We hope that the present contribution will facilitate experimental endeavours in molecular biology and spectroscopy to fully understand the role of retinol in transport, metabolism and signaling in carcinogenesis. It is reasonable to speculate that pharmacologically enhanced retinol activity may represent a method to inhibit certain types of cancer.

## Results

Here we will focus on information that can be extracted from Raman microscopy and Raman imaging of human central nervous system tissue from astrocytoma and medulloblastoma, as well as cell lines of normal human astrocytes (NHA), and glioblastoma (U-87 MG) cells. We will focus on cellular organelles such as the nucleus, lipid droplets, and mitochondria of NHA and U-87 MG cells.

## 1. Human cancer tissue

Fig 1 shows a typical Raman image and Raman spectra of central nervous system tissue from medulloblastoma and pilomyxoid astrocytoma compared with a white light microscopy image and MRI image. MRI provides the primary clinical information concerning the localization of the pathology, but Fig 1 shows that MRI has significantly limited spatial resolution in comparison to the Raman image. The Raman image offers unsurpassed spatial resolution combined with excellent spectral resolution. Therefore, Raman imaging is much more than simple microscopy because the Raman spectra embedded in the tissue image provide information about the biochemical composition of brain tissue with lipid-rich, protein-rich and mixed profiles, which are characteristic of brain white matter, tumours, and cortex with cellular, myelin-rich regions.[43,45–49]

**Fig 1.**
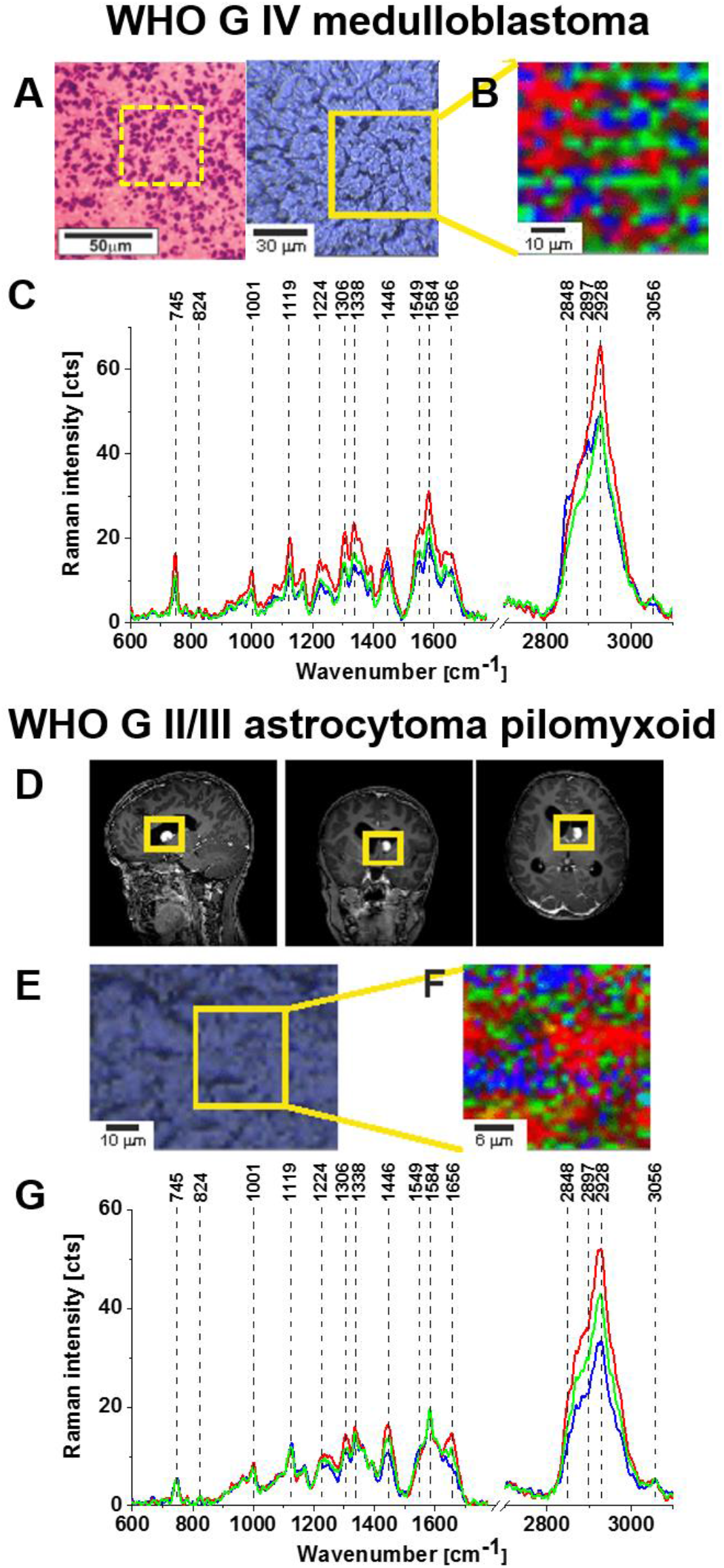
Raman spectroscopy analysis of medulloblastoma and astrocytoma. Microscopy image (A), Raman image (B), and characteristic Raman spectra (C) of WHO G IV medulloblastoma (P49). MRI images (D), microscopy image (E), Raman image (F) and characteristic Raman spectra (G) of WHO G II/III pilomyxoid astrocytoma (P51). The integration time for the Raman images was 0.5 s with a resolution step of 1 μm per 1 pixel for (F) and 3 μm per 1 pixel for (B), laser: 532 nm excitation, power: 10 mW. Size of map (B): 60×60 μm, number of Raman spectra 400; size of map (F): 30×30 μm, number of Raman spectra 900. The line colours of the spectra correspond to the colours of the Raman maps. The Raman maps were generated using basis analysis.

The red and blue areas in Fig 1 reflect the locations of proteins and lipids in specific brain regions, respectively. The Raman profiles of specific brain regions presented in Fig 1G reflect the different compositions of lipid-rich and protein-rich regions.

Numerous marked differences are apparent between the protein-rich and lipid-rich regions. The most significant differences are observed at 1584 cm^-1^ and 1656 cm^-1^.

To evaluate Raman imaging as a diagnostic tool for identifying tumours based on the biochemical composition of specific brain regions, one must better understand the differences presented in Fig 1. Thus, we compared the Raman spectra of the brain tumour tissue using different laser excitation wavelengths. This approach can reduce fluorescence at higher excitation wavelengths and might generate Raman resonance enhancement at lower excitation wavelengths for some tissue components that are non-visible for non-resonance conditions.

Fig 2 shows the Raman spectra of the high-grade medulloblastoma and low-grade astrocytoma tissues at an excitation wavelength of 785 nm, 532 nm, and 355 nm and different polarization configurations. The Raman spectra at different excitations are sensitive to the selective resonance Raman enhancement of the chromophores attached to the proteins and lipids. Therefore, from comparison of the Raman spectra at different excitation wavelengths one can learn about the modification of protein and lipid structures by the chromophore attached to the proteins or lipids that alter the Raman resonance conditions.

**Fig 2.**
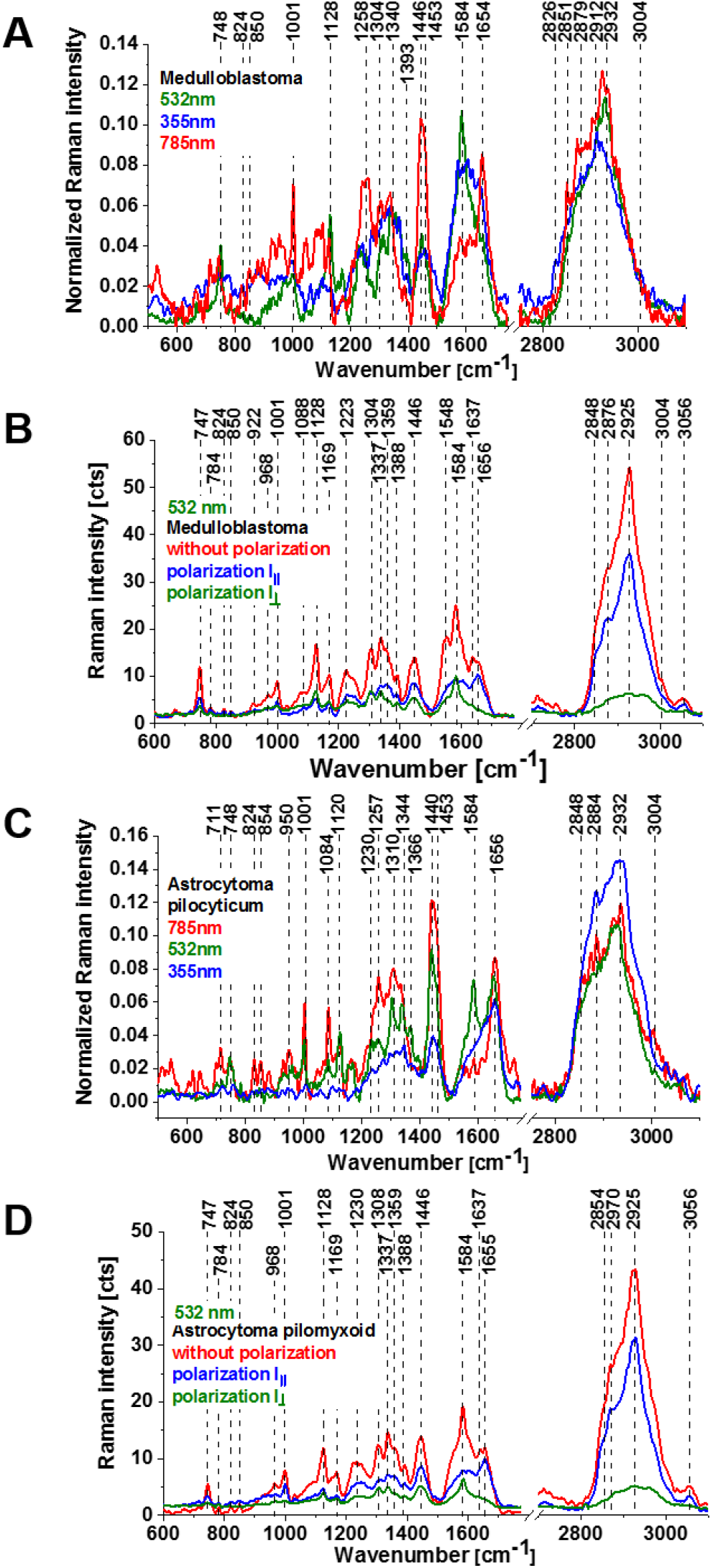
Raman vector normalized average spectra of human brain tissue from medulloblastoma (grade IV) (number of tissue samples n=11) (A), and astrocytoma (pilocyticum, grade I) (number of tissue samples n=8) (C) at 785 nm, 532 nm, and 355 nm wavelength excitation. Raman spectra for different polarization configurations at 532 nm: medulloblastoma (grade IV) (non-polarized Raman signal (red), parallel polarization I_II_ (blue), perpendicular polarization I_┴_ (green)) (B), and pilomyxoid astrocytoma (non-polarized Raman signal (red), parallel polarization I_II_ (blue), perpendicular polarization I_**┴**_ (green)) (D). Raman spectra in (B) and (D) were acquired at 0.5 s and 10 accumulations. Number of Raman spectra of (A) = 1600, (B) = 400 for each wavelength excitation, (C) = 1600, (D) = 600 for each wavelength excitation.

Additionally, polarized Raman spectroscopy can provide information about molecular symmetry and orientation as well as the symmetry of the bond vibrations, resulting in better visualization of conformational changes in cancerous and normal tissue structures. The different polarization configurations reflect the conformational modifications of the proteins and lipids.

Fig.2 shows dramatic differences due to the excitation as well as the polarization configurations in the Raman spectra for vibrations corresponding both to lipids and proteins. The most significant differences were observed at 1584 cm^-1^ and 1656 cm^-1^. Fig 2(A) shows that the Raman intensity of the band of amide I at 1656 cm^-1^ at 785 nm excitation increased in medulloblastoma compared with 355 and 532 nm. In astrocytoma pilocyticum (Fig 2 (C)) the amide I vibration intensity increased at 785 and 532 nm excitation compared with 355 nm. By contrast, the Raman intensity of the band at 1584 cm^-1^ increased at 355 nm and 532 nm and diminished at 785 nm, indicating that the chromophore (or chromophores) attached to the protein absorbs in the range from at 355 and 532 nm (Fig 2 (B,D)). The identification of the chromophores will be discussed in detail later in the paper.

The results obtained for different polarization configurations (Fig. 2 B and D) showed that the Raman signal at 1656 cm^-1^ reflecting vibrations of the α-helix protein structure became weaker under perpendicular polarization in comparison to the parallel polarization. By contrast, the Raman signal at 1584 cm^-^ was significantly enhanced under perpendicular polarization when compared to 1656 cm^-1^. The origin of the protein that is related to the band 1584 cm^-1^ will be discussed later in details, but the peaks ∼748, ∼1126, ∼1364 and ∼1584 cm^−1^ have been identified in pure cytochrome c and purified cytochrome c oxidase, one of the main proteins of the mitochondrial respiratory chain.[50,51] Furthermore, the peak ∼1364 and ∼1583 cm^−1^ were also associated with cytochrome c in isolated mitochondria from heart, liver and muscle.[50,51]. The peak at ∼1583 cm^-1^ was additionally identified in HeLa cells[52] and in yeast mitochondria.[53]

The other spectral regions with notable differences in the Raman signal in Fig. 2 A and C for various excitations were observed at approximately 1437-1444 cm^-1^ (fatty acids, triglycerides, CH_2_ or CH_3_ deformation modes[54,55]) and ∼2845-2940 cm^-1^ (CH_2_, CH_3_ stretching of lipids and proteins.[39,45,46,56]

Fig 2 shows the average Raman spectra providing information on lipids and proteins in human brain tissue of medulloblastoma and astrocytoma. To gain information about the chemical composition of specific organelles such as lipid droplets, the nucleus, mitochondria, and the cytoplasm inside the cells, we must focus our attention on individual single cells.

## 2. Single human normal and tumor cells

Fig 3 shows Raman images of glioblastoma cells (U-87 MG) compared with fluorescence images of dyes which specifically label different cellular organelles: Hoechst 33342, Oil Red O, and Mitotracker Red CMXROS labelling the nucleus (red), lipid droplets (blue), and mitochondria (magenta), respectively.

**Fig 3.**
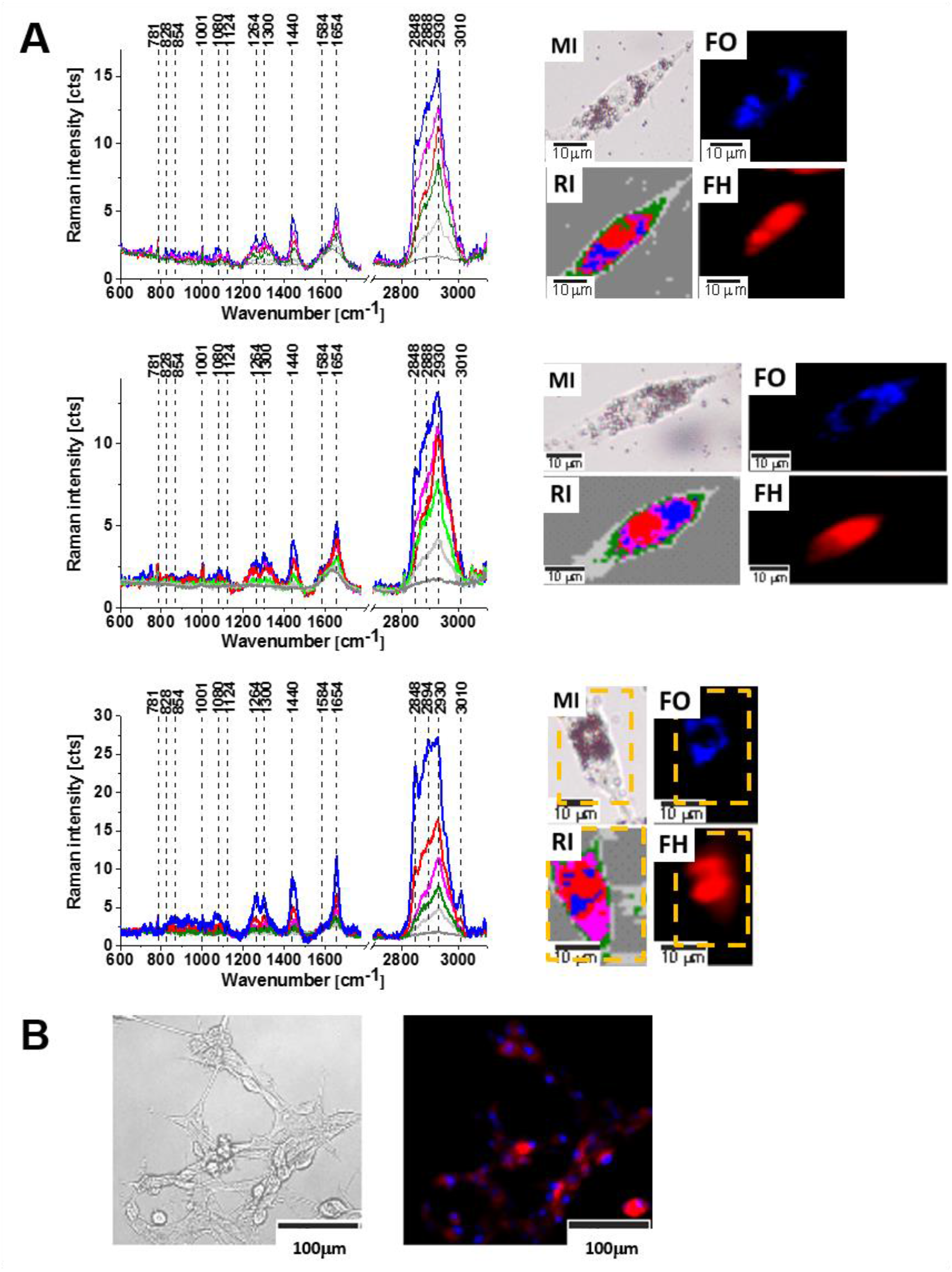
Confocal Raman spectroscopy analysis of the glioblastoma (U-87 MG) cells (number of cells n=3) at the 532 nm wavelength excitation. (A) Raman cluster spectra of nucleus (red), lipid droplets (blue), cytoplasm (green), mitochondria (magenta), cell border (light grey), area out of cell (dark grey), Raman cluster image (RI) (cell 1: 45×40 μm, cell 2: 55×30 μm, cell 3: 25×33 μm, resolution 1 μm, integration time 0.3 second, number of Raman spectra used for averaging 1800, 1650, 825, respectively), microscopy image after Oil Red O staining (MI), fluorescence image of Oil Red O staining (FO), and fluorescence image of Hoechst 33342 staining (FH). Microscopy and fluorescence image of U-87 MG cells stained with Mitotracker Red CMXROS and Hoechst 33342 (B).

The complete spectral and microscopic identification of organelles in normal and cancer cells allowed us to analyze the biochemical composition of lipid droplets, the nucleus and mitochondria by Raman mapping of specific cellular compartments.

First, we will focus on lipid droplets because Fig 3 MI (Oil Red O staining) and Fig 3 RI (Raman imaging) show a large concentration of lipid droplets around the nucleus. In our analysis we did not take into account the endoplasmic reticulum, which is also a lipid-rich structure, but the content of lipids is much lower than in LD consisting almost exclusively of lipids. We want to check whether this feature is characteristic for highly aggressive cancers such as glioblastoma or it is also typical for normal glial cells.

Fig 4 shows the microscope images, Raman images, and fluorescence images of normal human astrocytes (NHA) and high-grade glioblastoma cells (U-87 MG) compared with images of lipid droplets stained with Oil Red O dye.

**Fig 4.**
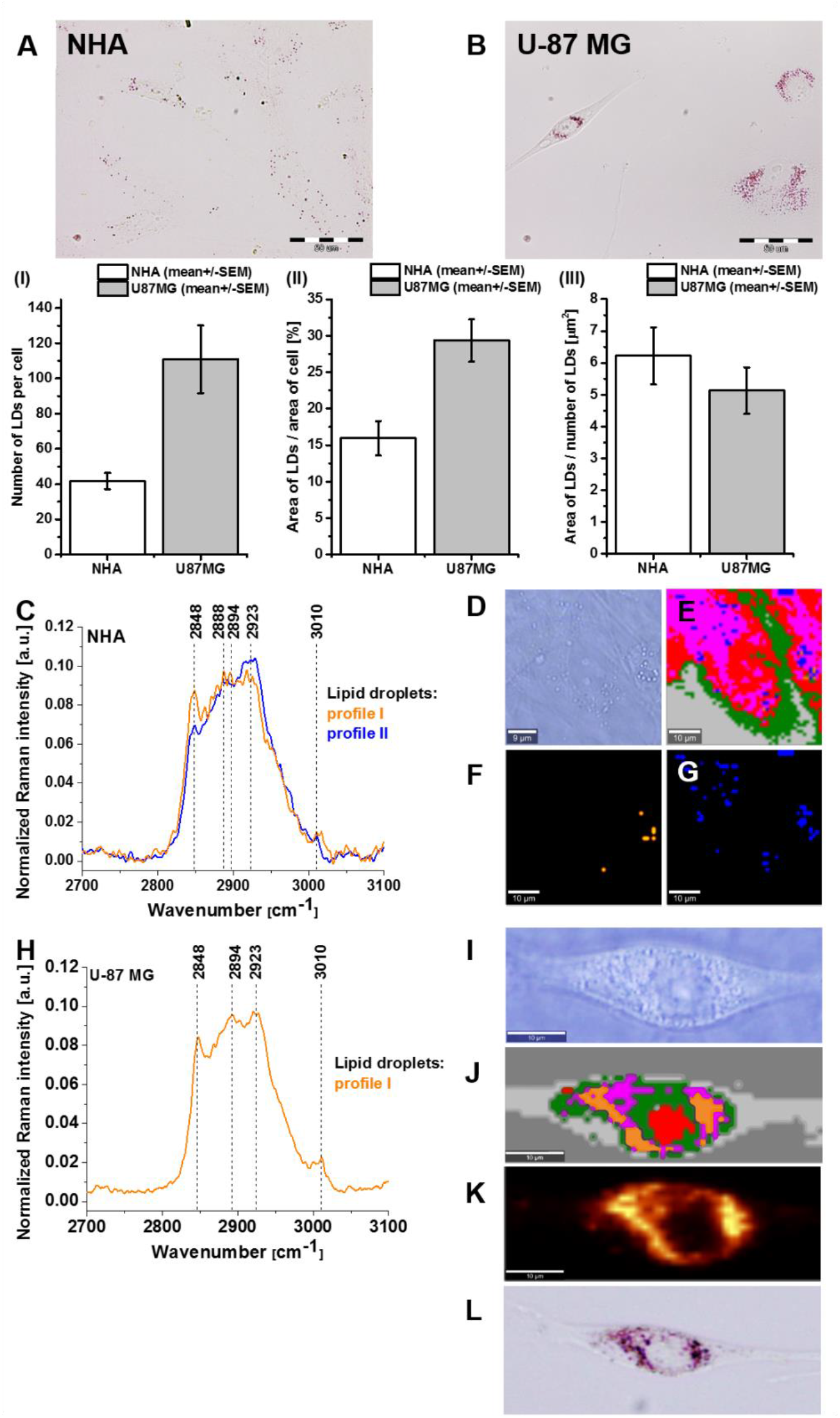
Comparison of lipid droplets in normal astrocytes (NHA) and high grade glioblastoma (U-87 MG) at the 532 nm wavelength excitation. Microscopy images of lipid droplets stained with Oil Red O dye in normal human astrocytes NHA (A) and high-grade glioblastoma U-87 MG (B and L) with bar plots of the total number of lipid droplets in NHA and U-87 MG (Fig.4 AB I), the area of LD per cell (Fig.4AB II) and the size of LDs were quantified by Image J software (Fig. 4 AB III). The results represent the means +/- SEM of at least 6 representative cells (n(NHA)=6, n(U-87 MG)=9), Raman spectra of lipid droplets of normal astrocytes, profile I – TAG (orange), profile II – retinyl esters (blue) (C), microscopy image (D), Raman cluster image (E)(image size 50×50 μm, resolution 1 μm, integration time 0.3 second, number of Raman spectra 2500) and Raman image of the distribution of lipid droplets with profile I (F) and profile II (G) in NHA. Average Raman spectrum of lipid droplets obtained from cluster analysis (H), microscopy image (I), Raman cluster image (J)(image size 55×20 μm, resolution 1 μm, integration time 0.3 second, number of Raman spectra 1100) and fluorescence image of Oil Red O (K) and microscopy images of Oil Red O-stained lipid droplets (L) of U-87 MG high-grade glioblastoma. Raman analysis were performed at 532 nm laser excitation. Nucleus was labeled by red, lipid droplets – blue/orange, cytoplasm - green, mitochondria - magenta, cell border - light grey and area out of cell - dark grey.

Detailed inspection of Fig 4 provides valuable information related to the number of lipid droplets and their chemical compositions. First, Fig 4 (A,B,L) clearly shows that cancer cells of U-87 MG high-grade brain glioblastoma contain significantly more lipid droplets than NHA normal astrocyte cells. To quantify the number of lipid droplets in normal astrocytes and high-grade glioblastoma, we counted the number of lipid droplets per cell in the microscopy images of Oil Red O-stained cells and Raman images of fixed regions. Fig. 4 AB (I, II, III) shows the bar plots of the total number of lipid droplets in NHA and U-87 MG, the area of LD per cell and the size of LDs.

U-87 MG cell line contains the average number of lipid droplets of 110 per cell and 41 LDs/cell for NHA, respectively (Fig. 4AB I). The number of lipid droplets in individual cells was also scored quantitatively by cluster analysis by measuring the area of separated clusters corresponding to lipid droplets, mitochondria and nucleus. (Fig. 4 AB II). The results and conclusions from Oil Red O-stained cells and Raman images of fixed regions were identical. It is interesting to notice that the number of lipid droplets in normal cells NHA is smaller than in cancer cells of U-87 MG, but the size of the single lipid droplet is larger (Fig. 4 AB III).

The increased number of cytoplasmic lipid droplets in human brain cells must be closely related to the increased rate of de novo lipid synthesis in cancer cells; An increased number of lipid droplets was correlated with increased aggressiveness of the brain cancer.[57,58] However, the detailed mechanisms are still unknown.

This feature of enhanced de novo lipid synthesis seems to be universal because recently, we have shown that the increased number of lipid droplets is also typical for breast cancer and can be considered as a hallmark of cancer aggressiveness.[45]

Second important feature that can be observed from the results presented in Fig. 4 is related to the chemical composition of lipid droplets in normal and tumorous cells.

Based on a detailed analysis of the chemical composition of lipid droplets of NHA normal astrocytes and U-87 MG high-grade glioblastoma cells by combining cluster analysis and Raman imaging, we found two types of lipid droplet Raman profiles, which are presented in Fig 4(C and H). The composition of lipid droplets in normal droplets represented by Profile I. Both profiles characterize the vibrations in the high frequency region 2700-3100 cm^-1^ and correspond to the polyene chains of retinoids as well as chains of hydrocarbons of lipids that are easily visible in non-resonant Raman conditions. Detailed analysis in our previous papers showed that the profile I represents lipid droplets filled with triglycerides (TAG).[45] The identification of the Profile II will be discussed in detail later in the paper where we will show that it represents lipid droplets filled with retinoids.

Detailed inspection into Fig. 4 F and 4 G clearly shows that normal astrocytes NHA are dominated by the profile II. In contrast, U-87 MG high-grade glioblastoma is dominated exclusively by lipid droplets filled with triglycerides (TAG) (Profile I). Indeed, the comparison between the Raman images of the distribution of retoinoid - rich lipid droplets in Fig 4(G) with TAG-rich lipid droplets in Fig 4(F) provides clear evidence that most lipid droplets in normal astrocytes NHA accumulate retoinoid - rich lipid droplets. By contrast, lipid droplets in glioblastoma almost exclusively accumulate TAG-rich lipid droplets (Fig 4(J and H)). As it is well known, TAGs in lipid droplets are related mainly to their function as energy storage deposits.[46] The role of retoinoid - rich lipid droplets is still unknown, but recent research has revealed exciting new aspects of retinoids importance in the regulation of lipid metabolism, cell signaling, membrane trafficking and control of the synthesis and secretion of inflammatory mediators.[59] In the view of the presented results we will now concentrate on these potential functions of lipid droplets, which are related to alterations in accumulation of retinoids in lipid droplets.

To contribute to this aspect of retinoids importance we will use Raman imaging to examine in detail differences in chemical composition of lipid droplets, mitochondria and nucleus in normal astrocytes and high-grade malignant human brain cells (glioblastoma). To examine role of retinoids we will use various wavelengths of excitation in Raman spectroscopy, because the major chemical components in cell organelles can be selectively monitored by such resonance Raman enhancement effect and can be clearly visualized by Raman imaging at physiological concentrations.

To define the Raman resonance conditions we must learn about the absorption spectra of the main biochemical components identified in tissues and single cells.

Fig 5 shows the absorption spectra of retinoids, TAG (glyceryl tripalmitate) and retinol binding protein I (RBP-1) and cytochrome c. One can see that TAG and RBP-1 exhibit negligible absorption in the spectral range of 200-800 nm. In contrast, retinoids show significant absorption abound 330-388 nm. Therefore, the excitation with 355 nm will reveal vibrations of retinoids at physiological concentrations due to Raman resonance enhancement that are non-visible for non-resonance conditions at 532 nm and 785 nm.

**Fig 5.**
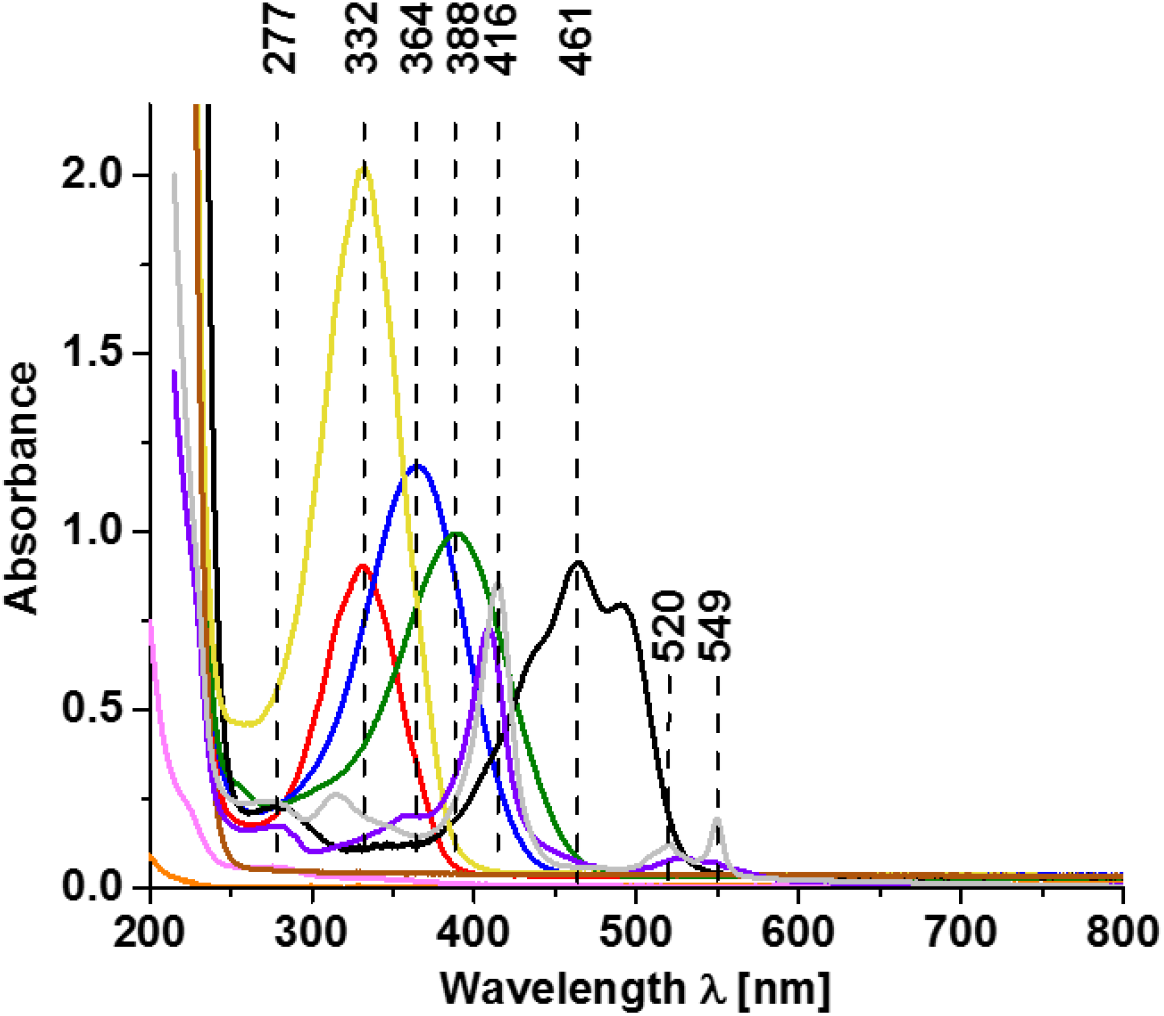
Absorption spectra of retinoids, TAG (glyceryl tripalmitate), cytochrome c and retinol binding protein I. Retinol (c=0.2 mM in chloroform, red), retinoic acid (c=0.2 mM in chloroform, blue), all-trans retinal (c=0.1 mM in chloroform, green), β-carotene (c=0.3 mM in chloroform, black), retinyl palmitate (c=0.2 mM in chloroform, yellow), glyceryl tripalmitate (1 mM in hexane, orange), retinol binding protein I (1.5 mM, magenta), cytochrome c (0.18 mM in PBS; Fe^3+^ (grey), Fe^2+^ (violet)) and chloroform (brown), with a cuvette path length of 1 mm.

## 3. Retinoids

To address questions regarding selectively enhanced effects on lipid droplet alterations upon laser excitation, we examined first the resonance and non-resonance Raman effects in model systems of TAGs, RBP-1 and retinoids. The results are presented in Fig 6. Figs 6(A) and 6(B), show the Raman spectra of retinyl palmitate (RP) in oleic acid (OA) at 532 nm and 355 nm, respectively, at different polarization configurations.

**Fig 6.**
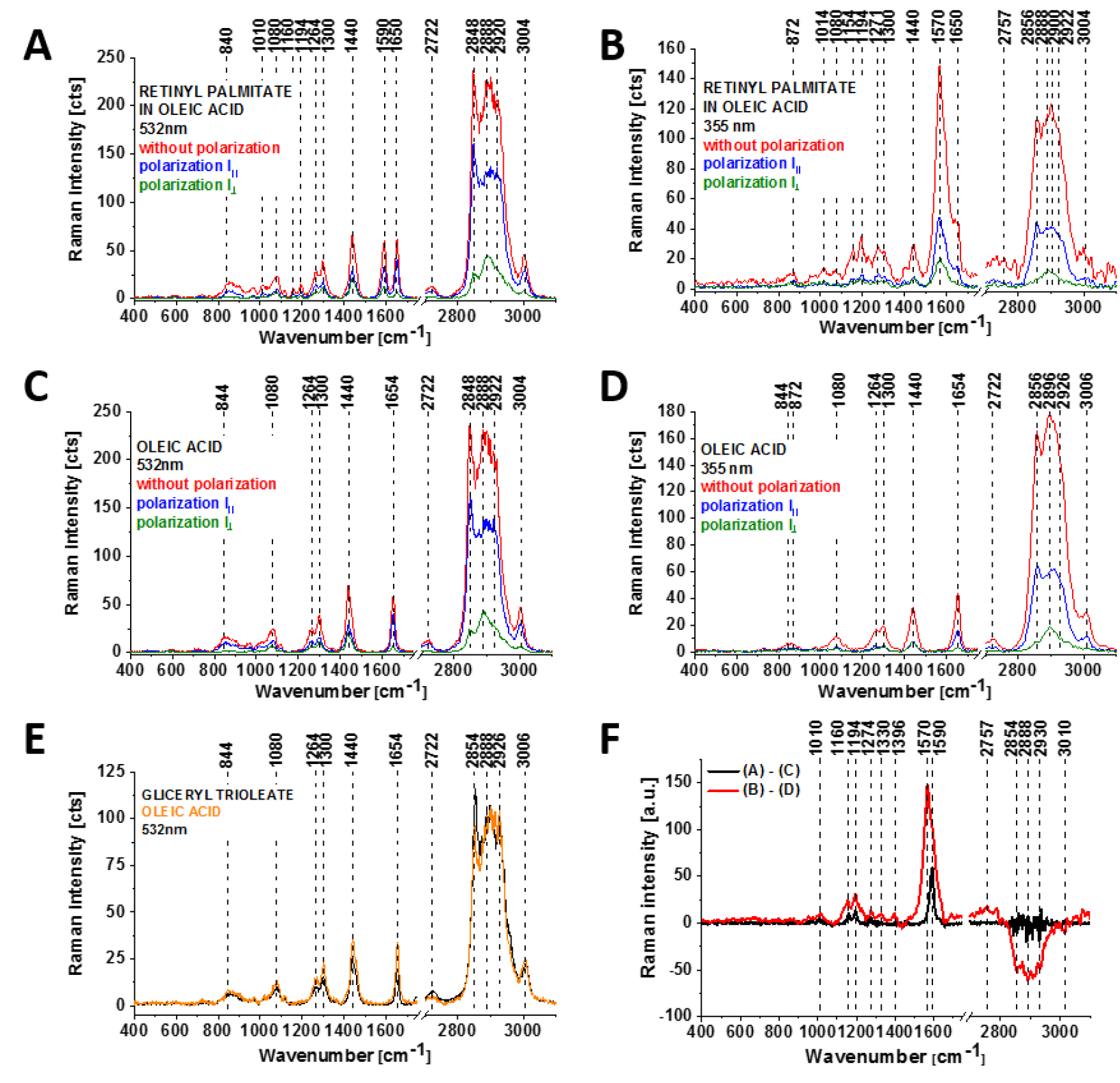
Raman spectra of retinyl palmitate (RP) in oleic acid (OA) (A,B) and oleic acid. (C,D) under different polarization configurations (non-polarized (red), parallel polarization III (blue), perpendicular polarization I┴ (green) at 532 nm (A,C) and 355 nm (B,D) and Raman spectra of TAG (glyceryl trioleate) and oleic acid (E) at 532 nm. (F) Difference spectra of (A)-(C) and (B)-D).

The results presented in Fig 6 clearly demonstrate that TAG compounds (OA (Fig 6 (C,D), glyceryl trioleate (E)) show no Raman resonance enhancement in contrast to retinyl palmitate (Fig 6 (A,B)), with the characteristic band of RP at 1590 cm^-1^ for 532 nm excitation. This band disappears at 355 nm excitation, as presented in Fig 6B and a new band at 1570 cm^-1^ corresponding to all-trans retinal and/or retinoic acid appears as a result of photochemical reaction presented in Fig. 7.[60] The Raman band at 1570 cm^-1^ is significantly enhanced in Fig. 6 B at 355 nm by the resonance Raman effect due to the absorption of all-trans retinal and retinoic acid at 388 nm and 364 nm as presented in Fig 5.

**Fig 7.**
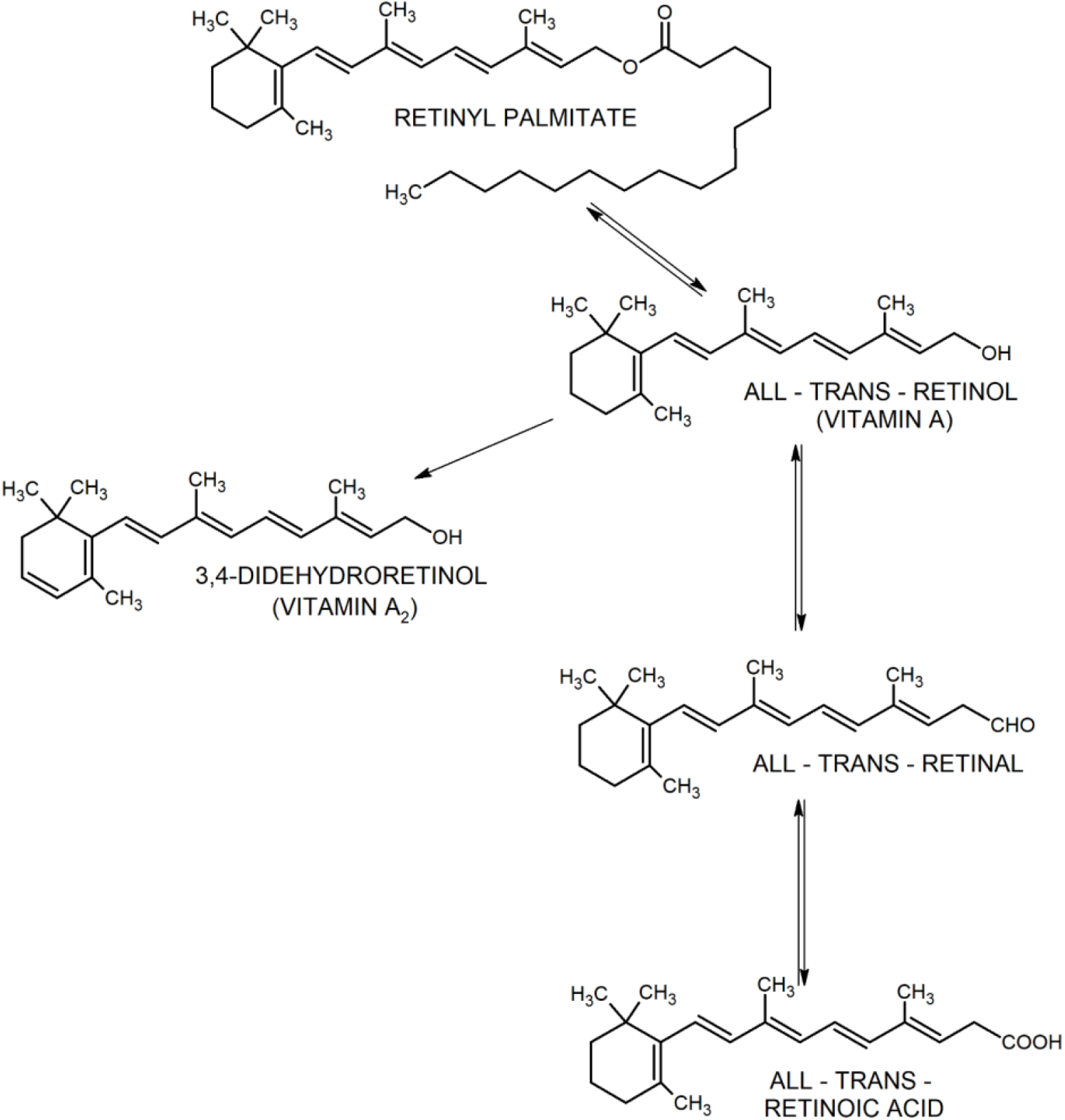
Photochemical and enzymatic reactions of selected retinoids upon 355 nm excitation.[60].

The results presented herein for model systems demonstrate that Raman microscopy and imaging are excellent tools to monitor the role of retinoids in single cells and tissues.

Fig 8 (A, B) shows the Raman spectra of retinoids: retionol, retinoic acid, all-trans retinal, retinyl palmitate, with the characteristic bands at 1570 cm^-1^ (retinoic acid, all-trans retinal) and 1590 cm^-1^ (retinol, retinyl palmitate). The high frequency vibrations in the range 2800-3100 cm^-1^ show three types of Raman band profiles: profile A – for retinol, all-trans retinal, RBP-1; profile B - retinoic acid; profile C - retinyl palmitate. Detailed analysis of biochemical profiles showed that Profile II in Fig 4 and profile A in Fig 8 are identical suggesting that the lipid droplets in normal astrocytes NHA are filled with retinol and/or retinyl esters.

**Fig 8.**
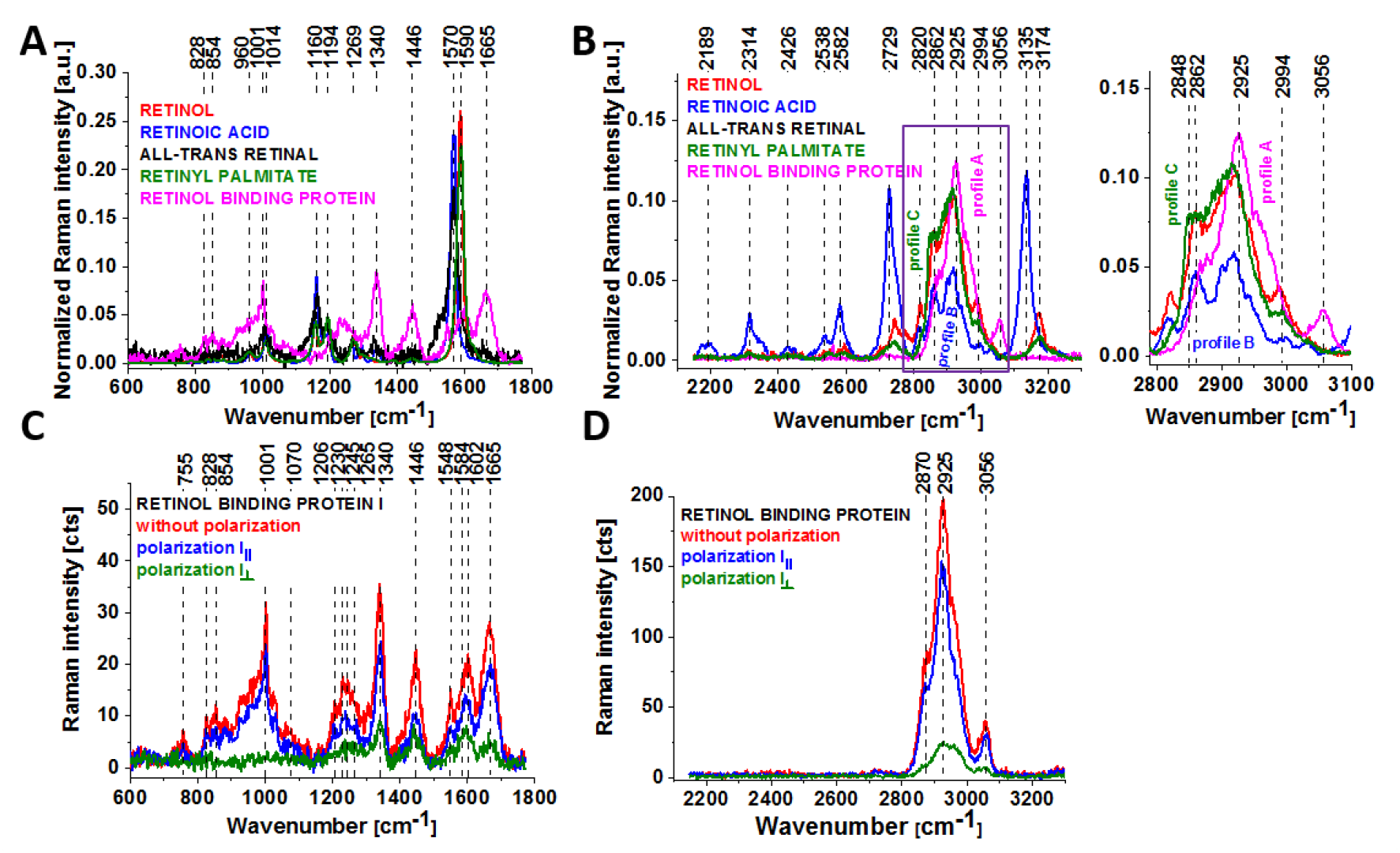
Raman spectra of the retinoids (A,B) and non-polarized and polarized Raman spectra of retinol binding protein RBP-1 (C,D) (non-polarized (red), parallel polarization (blue), perpendicular polarization (green)). Raman spectra were recorded at 532 nm wavelength excitation.

Fig 8 (C) shows the Raman spectra of retinol-binding protein (RBP-1) with the characteristic vibrations of amide I (1665 cm^-1^), amide II (1548 cm^-1^), tyrosine (1603 cm^-1^), tyrosine doublet at 828 cm^-1^ and 854 cm^-1^ representing the tyrosine residue of RBP-1 corresponding to a Fermi-resonance between the first overtone of the aromatic out-of-plane ring bending mode and the breathing fundamental aromatic ring[61,62], and the band at 1265 cm^-1^ corresponding to amide III and red-shifted to the 1230 cm^-1^ band of amide III upon phosphorylation.[62,63] Vibrations of the PO_4_- phosphate group of phosphorylated tyrosine were observed at 1070 cm^-1^ and corresponded to the O-P-O symmetric stretching mode.[62–66]

## 4. The resonance and non-resonance effects of retinoids in the biochemical Raman maps of cellular organelles in cell cultures

We found that retinoids in Fig 6 are sensitive to the resonance Raman enhancement. Now, we want to find and examine the resonance and non-resonance effects of retinoids in the biochemical Raman maps of cellular organelles in cell cultures of normal astrocytes (NHA) and glioblastoma (U-87 MG) as a function of the laser wavelength excitation to examine the Raman signal under resonance and non-resonance conditions.

Fig 9 shows the Raman spectra and images of U-87 MG glioblastoma and NHA normal astrocytes at 355, 532 and 785nm excitations.

**Fig 9.**
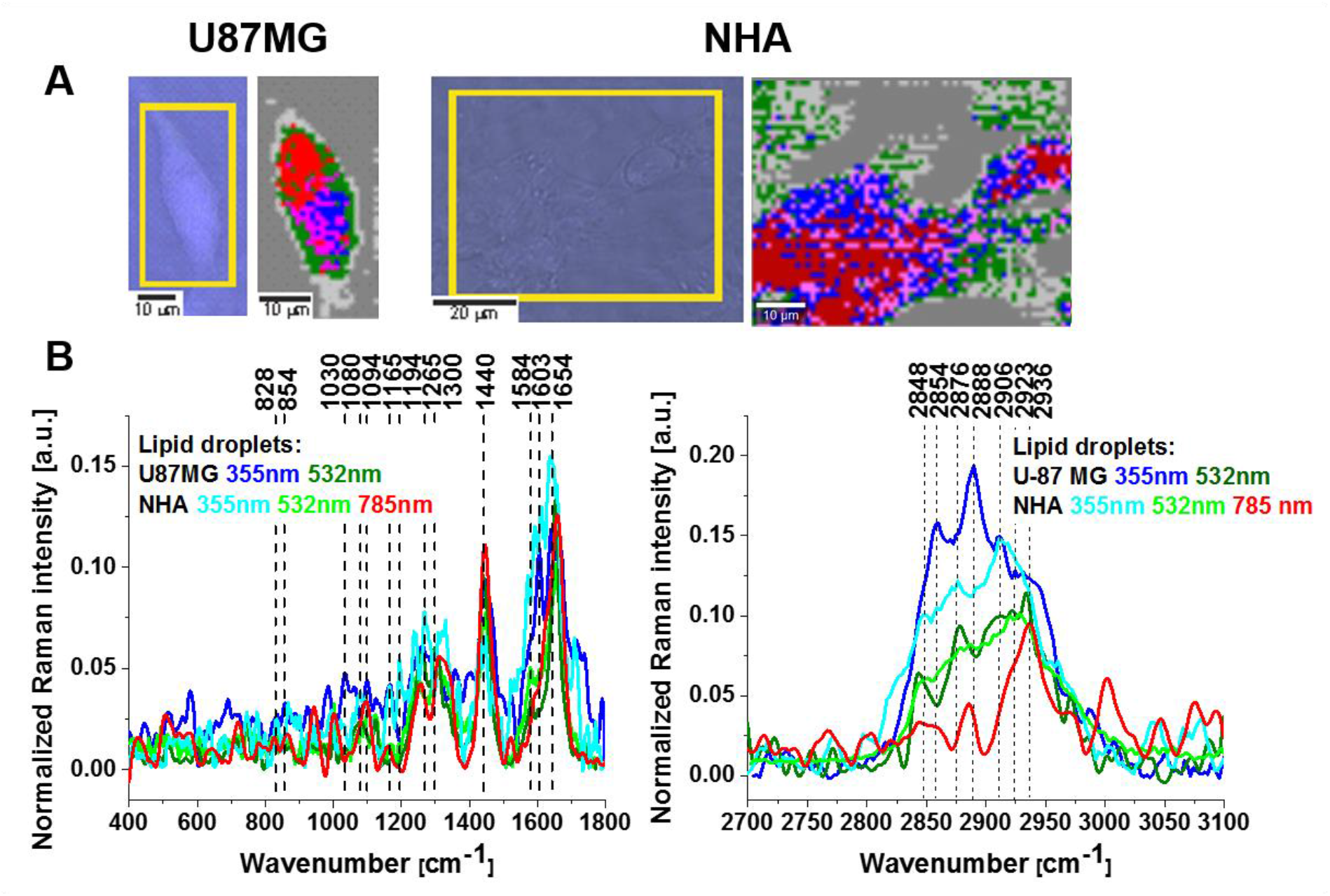
Comparison of normal astrocytes (NHA) and high grade glioblastoma (U-87 MG) at the 355 nm wavelength excitation. Microscopy images, Raman images (excitation at 355 nm, image size of U-87 MG: 20×45 μm, image size of NHA: 65×50 μm, resolution 1 μm, integration time 0.3 second, number Raman spectra used for averaging 900 (U-87 MG) and 3250 (NHA)) (A) and Raman spectra of lipid droplets at 355, 532 and 785 nm (B).

The red, blue and pink areas in Raman cluster images in Fig 9(A) represent nuclei, lipid droplets and mitochondria, respectively. Lipid droplets are typically localized in strings adjacent to the nucleus (as seen in Fig 3) or randomly distributed in the cytosol, the mitochondria are clustered around the lipids. It has been suggested that lipid droplets are not only an energy storage source for mitochondria, but also mitochondria play an important role in helping to build the lipid droplets.[67]

A detailed inspection of Fig 9(B) reveals notable differences in the Raman vibrations of lipid droplets at different excitation wavelengths. Fig 9(B) shows that the average Raman profile of lipid droplets is significantly different at different excitations: 355 nm, 532 nm and 785 nm corresponding to resonance and non-resonance conditions both for cancer cells U-87 MG and normal cells NHA. Fig 9 B shows that the Raman intensities of the bands at 1584 cm^-1^ (cytochrome c) and 1603 cm^-1^ (the tyrosine residue of RBP-1) increased at 355 nm excitation and diminished at 532 nm and 785 nm, indicating that the chromophore attached to the proteins (cytochrome c, the tyrosine residue of RBP-1) absorbs in the range of 355 nm corresponding to absorption of retinoids (see Fig. 5). Moreover, the profile in the high frequency region at 355 nm for U-87 MG can be easily identified as the vibration retinyl palmitate chain whereas the profile for NHA corresponds to retinol. The higher resonance Raman enhancement observed in Fig. 9 B in the high frequency region for U-87 MG droplets (filled with retinyl palmitate) than for NHA lipid droplets (filled with retinol) corresponds well with the resonance enhancement conditions in Fig. 5.

To summarize, the results for the brain single cells are consistent with those obtained for the brain tissues presented in Fig 2, indicating that both lipid droplets in single cells and tissues contain compounds that are resonance Raman enhanced at 355 nm excitation corresponding to Raman resonance conditions of retinoids. Therefore, Fig. 9A presents the distribution of retoinoid -rich lipid droplets (blue color) in normal astrocytes (NHA) and high grade glioblastoma (U-87 MG). It is worth of noticing that the normal NHA cells contain much more retinoids than the high grade glioblastoma (U-87 MG) cells. In contrast, U-87 MG contain more TAGs than the normal cells NHA as we presented in Fig. 4.

The results presented thus far provide strong evidence that lipid droplets accumulate retinoids and TAG lipids and the relative amount of retinoids/TAG lipids may be correlated with cancer aggressiveness. More aggressive cancer cells accumulate TAG lipids.

As has been suggested that lipid droplets are not only an energy storage source for mitochondria, but also mitochondria play an important role in helping to build the lipid droplets [67] we next detailed the differences in the chemical composition of lipid droplets, mitochondria and the nucleus in normal astrocytes and high-grade malignant human brain cells (glioblastoma). We compared the alterations in the chemical composition of lipid droplets, nuclei and mitochondria of normal astrocytes NHA and high-grade glioblastoma (U-87 MG).

Fig 10 shows a comparison of the average Raman spectra (normalized by vector norm) obtained from the cluster analysis of lipid droplets, mitochondria and nuclei of normal human astrocytes (NHA) and high-grade glioblastoma (U-87 MG) at 532 nm excitation.

**Fig 10.**
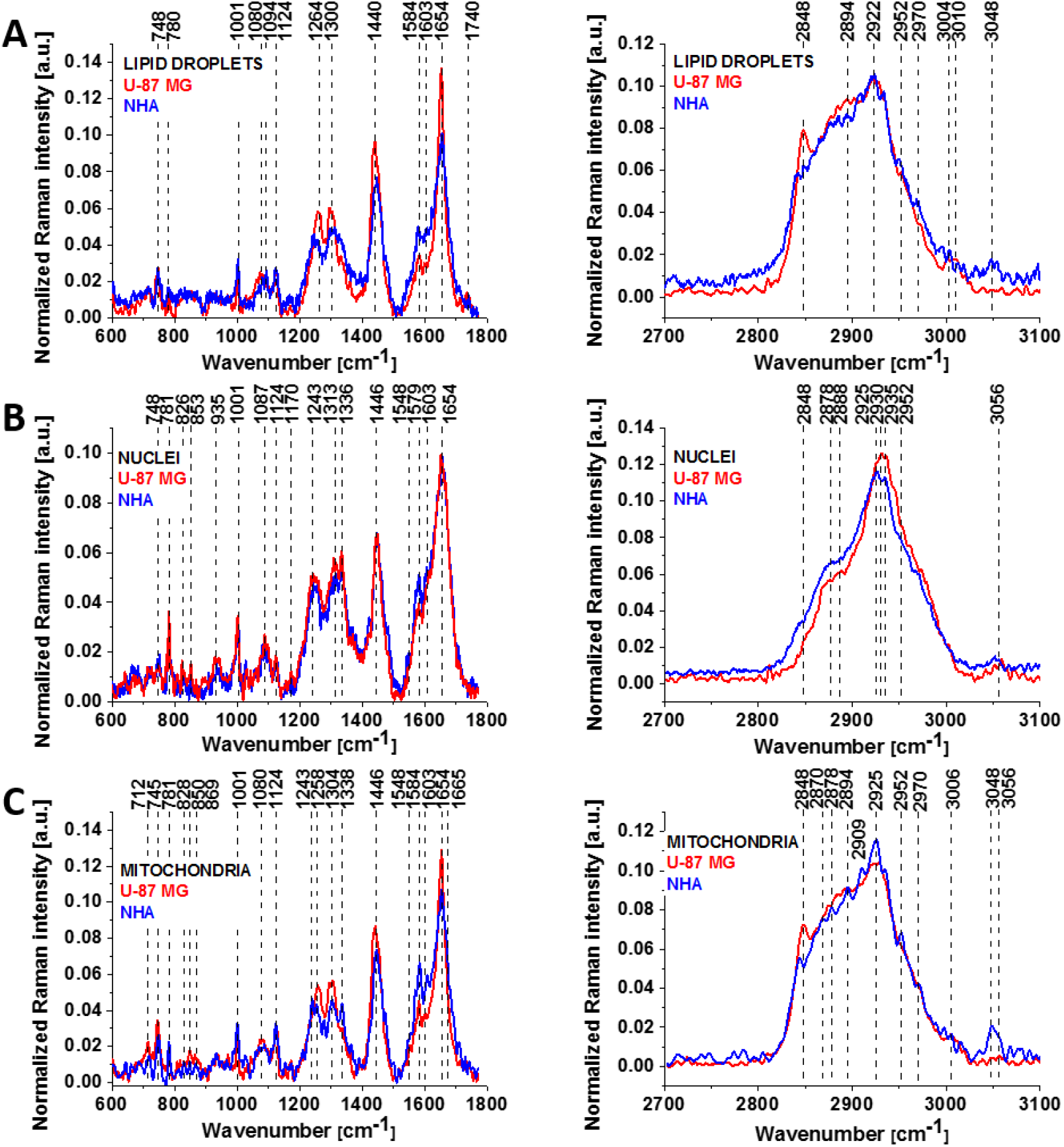
The average Raman spectra (normalized by vector norm) obtained at 532 nm wavelength laser excitation from the cluster analysis of lipid droplets (A), nuclei (B) and mitochondria (C) of normal human astrocytes (NHA, blue) and high-grade glioblastoma (U-87 MG, red). Number of cells n(NHA)=4, n(U-87 MG)=3; number of NHA and U-87 MG Raman spectra used for averaging 4725 and 4275, respectively.

Detailed inspection of Fig 10 revealed that clear differences between NHA normal human astrocytes and U-87 MG high-grade glioblastoma in all studied organelles occurred for the same vibrations that were sensitive to Raman resonance enhancement due to the retinoid absorption at 355 nm excitation. They are observed at 748 cm^-1^, 1584 cm^-1^ (cytochrome c), 828 cm^-1^ and 854 cm^-1^, 1603 cm^-1^ (tyrosine residue of RBP-1), 1654 cm^-1^ and in the high frequency region at 2848 cm^-1^ (corresponding to TAG lipids) and 2923 cm^-1^ corresponding to both TAG, retinoids, and retinol binding proteins. Retinol binding proteins belong to lipocalin family with the concentration of RBPs in serum around 40 to 50 μg/ml, which indicates that they are undetectable at non-resonance Raman conditions at 532 nm.

They are detectable only in the complex RBP-retinol [68] at 355 nm excitation due to retinol Raman enhancement which we observed in Fig. 9. In contrast, the cytochrome family with Q-band absorption at 500-550 nm is in resonance with 532 nm excitation and the band 1584 cm^-1^ may correspond to cytochrome c.[58]

Fig 10 shows that the Raman signal intensity of the band at 1584 cm^-1^ in lipid droplets is significantly enhanced for normal astrocyte (NHA) compared with high-grade glioblastoma (U-87 MG). The 1584/1654 Raman intensity ratio of the bands at 1584 cm^-1^ and 1654 cm^-1^ was 0.61 for lipid droplets in NHA and 0.25 for lipid droplets in U-87 MG. Moreover, the 2848/2923 Raman intensity ratio of the bands at 2848 cm^-1^ and 2923 cm^-1^ was 0.48 for NHA and 0.80 for U-87 MG, indicating a lower concentration of TAG lipids (and a higher concentration of retinyl esters) in NHA normal cells than U-87 MG cancer cells. The 1584/1654 Raman intensity ratios for mitochondria were also significantly higher (0.65) for NHA than U-87 MG (0.35), indicating a higher concentration of retinol binding proteins in the mitochondria of normal NHA cells. The 2848/2923 ratio for mitochondria was 0.48 in NHA and 0.70 in U-87 MG, indicating a lower concentration of TAG lipids in NHA normal cells than U-87 MG cancer cells. The 1584/1654 Raman intensity ratio for nuclei was also significantly higher (0.50) for NHA than U-87 MG (0.38).

The results presented in Fig 10 and Table 1 reveal the lower expression of proteins represented by the band at 1584 cm^-1^ in human brain cancer. Second, the Raman signal intensity of the band at 2848 cm^-1^ in lipid droplets was significantly higher for high-grade cancer of glioblastoma (U-87 MG) than for normal astrocytes (NHA), indicating a higher concentration of TAG in lipid droplets. The higher concentration of TAG was related to the increased amount of cytoplasmic lipid droplets in human glioblastoma cells in comparison to normal astrocytes. By contrast, the intensity of the band at 2848 cm^-1^ in the nucleus was higher in NHA than in U-87 MG.

**Table 1.**
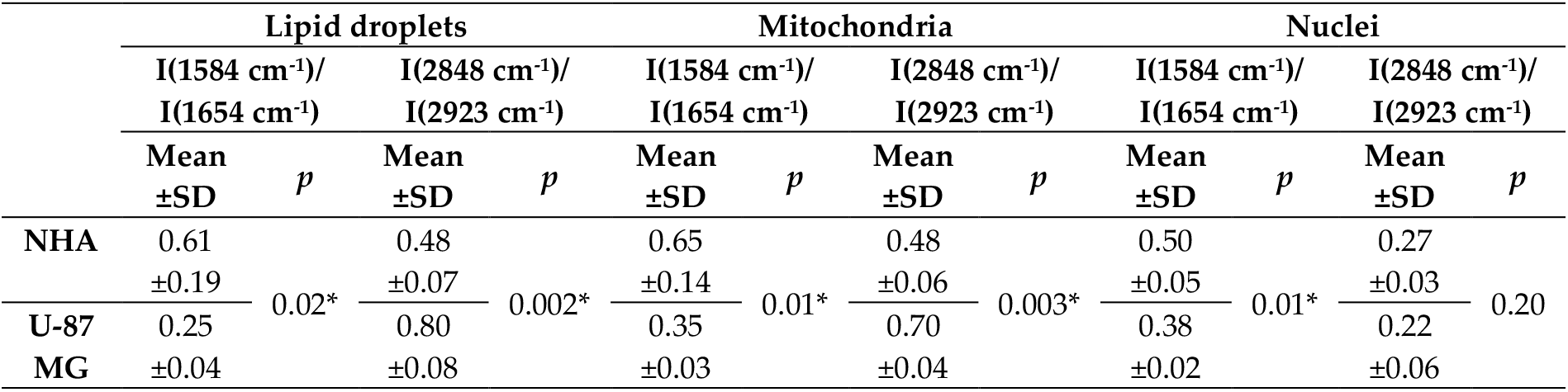
Comparison of the Raman intensity ratio of lipid droplets, mitochondria and nuclei for NHA and U-87 MG cell lines, presented as the mean±SD, SD - standard deviation, *p* – probability value (n(NHA)=4, n(U-87 MG)=3). *, where p<0.05. Raman bands intensity were taken from normalized by vector norm spectra.

|  | Lipid droplets |  |  |  | Mitochondria |  |  |  | Nuclei |  |  |  |
| --- | --- | --- | --- | --- | --- | --- | --- | --- | --- | --- | --- | --- |
| | I(1584 $\text{cm}^{-1}$)/<br>I(1654 $\text{cm}^{-1}$) | | I(2848 $\text{cm}^{-1}$)/<br>I(2923 $\text{cm}^{-1}$) | | I(1584 $\text{cm}^{-1}$)/<br>I(1654 $\text{cm}^{-1}$) | | I(2848 $\text{cm}^{-1}$)/<br>I(2923 $\text{cm}^{-1}$) | | I(1584 $\text{cm}^{-1}$)/<br>I(1654 $\text{cm}^{-1}$) | | I(2848 $\text{cm}^{-1}$)/<br>I(2923 $\text{cm}^{-1}$) | |
| | Mean<br>±SD | $p$ | Mean<br>±SD | $p$ | Mean<br>±SD | $p$ | Mean<br>±SD | $p$ | Mean<br>±SD | $p$ | Mean<br>±SD | $p$ |
| NHA | 0.61<br>±0.19 | 0.02* | 0.48<br>±0.07 | 0.002* | 0.65<br>±0.14 | 0.01* | 0.48<br>±0.06 | 0.003* | 0.50<br>±0.05 | 0.01* | 0.27<br>±0.03 | 0.20 |
| U-87<br>MG | 0.25<br>±0.04 | | 0.80<br>±0.08 | | 0.35<br>±0.03 | | 0.70<br>±0.04 | | 0.38<br>±0.02 | | 0.22<br>±0.06 | |

## 5. Single human tumor cells incubated with retinol

To clarify better the origin of the band at 1584 cm^-1^ assigned to cytochrome c, which is crucial for understanding the role of retinoids in cancer we incubated the glioblastoma cells (U-87 MG) with retinol (ROL). Fig 11 shows the Raman spectra of nucleus, lipid droplets and mitochondria in the U-87 MG cells incubated with ROL. As one can see from Fig 11 ROL (spectrum marked in blue) exerts a spectacular effect on the Raman spectra of glioblastoma cells, particularly on the band at 1584 cm^-1^ when compared with the control cells. First, the maximum of the Raman band observed at 1584 cm^-1^ for control cells is shifted to 1590 cm^-1^ for cells incubated with ROL. We showed that the vibration at 1590 cm^-1^ corresponds to the vibrations of retinol (see Fig 8A) indicating that the lipid droplets and mitochondria are filled with retinol when incubated with ROL. In contrast to lipid droplets and mitochondria (Fig. 11 A and C), ROL has lower influence on nuclei (Fig.11B).

**Fig 11.**
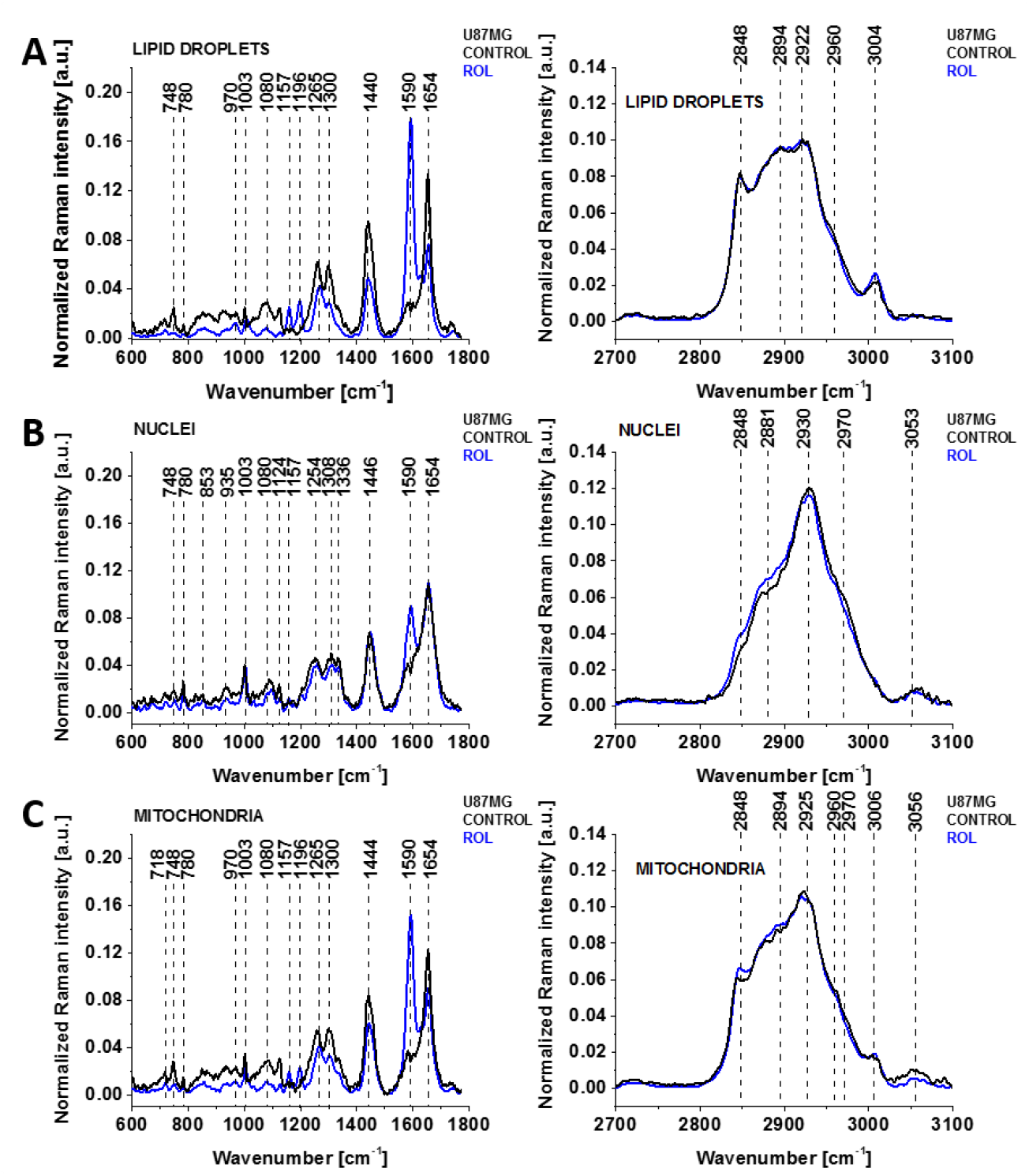
The average Raman spectra (normalized by vector norm) obtained at 532 nm wavelength laser excitation from the cluster analysis of lipid droplets (A), nuclei (B) and mitochondria (C) of high-grade glioblastoma (U-87 MG) incubated with retinol (ROL, blue). Concentration of 1 μM, incubation time of 24 hours. Number of cells n(U-87 MG (control))=3, n(U-87 MG (ROL))=3; number of U-87 MG (control) and U-87 MG (ROL) Raman spectra used for averaging 4275 and 9825, respectively.

In the view of the results for normal and high-grade tumor of glioblastoma it is exciting to compare the results from Fig. 11 with Raman spectra of human brain tissue from medulloblastoma (grade IV) and astrocytoma (pilocyticum, grade I) at 532 nm from Fig. 2. One can see that the Raman band at 1584 cm^-1^ is very strong both in the cancer cells when supplemented with ROL and in cancer tissue at 532 nm excitation. In contrast, we showed that the band at 1584 cm^-1^ in the normal brain cells (NHA) is much weaker (Fig. 10). We want to stress that the retinol concentration of 1 μM cannot be observed at non-resonance Raman conditions at 532 nm excitation. It indicates that retinol must be coupled to a protein that has absorption at around 532 nm to create Raman resonance enhancement. As can be seen from Fig. 5 RBP-1 does not exhibit absorption at 532 nm in contrast to cytochrome c. It suggests that retinoids are coupled to cytochrome c, which may modulate electrostatic gradient in the mitochondrial cell membrane.

## Discussion

The results presented in this paper for brain tissues and cells fully support the importance of retinoids in brain carcinogenesis.

Based on detailed Raman imaging analysis of lipid droplets, mitochondria and nucleus, we identified a protein that exhibits an enhanced Raman resonance signal at 1584 cm^-1^, which is shifted to 1590 cm^-1^ when incubated with retinol when excited with 532 nm corresponding to the absorption of cytochrome c. The 1584 cm^-1^ signal is particularly strong in lipid droplets and mitochondria. We assigned the Raman signal at 1584 cm^-1^ to a protein that has β sheets structure and vibrational signatures characteristic of the cytochrome c and phosphorylation activity of tyrosine residues of RBP-1.[39,58] The resonance Raman enhancement at 355 nm indicates that retinoids, particularly retinol, are coupled with the RBP-1 protein. On the other side, the resonance Raman enhancement of retinol at 1590 cm^-1^ for laser excitation of 532 nm provide evidence that the protein is coupled with retinoids. We suggest that the 1584 cm^-1^ band originates from cytochrome c involved as an intermediate in the electron transport chain in mitochondria, where the electron transport is blocked due to cancer development. Cytochrome c plays a key role in the respiratory activity in mitochondria and hence is essential for the life of a normal and tumor cells.[58]

Our results obtained in this paper are consistent with data obtained for retinol binding protein I (RBP-1)[1] showing that wild-type RBP-1 (also known as CRBP-1) has lower expression in cancers. The results presented in Fig 10 and Table 1 reveal the lower expression of the proteins represented by the band at 1584 cm^-1^ in human brain cancer. Our results are also consistent with those reported recently.[1,27,33] where lower expression of cellular retinol-binding protein (CRBP-1) has been observed in breast and other human cancers.[1,27,33] Recent research has demonstrated that lower expression of CRBP-1 decreases the ability of RBP-retinol complex to phosphorylate STRA6 and, later, JAK2 and STAT5.[2]

The results presented in this paper and analyzed in the context of recent studies[1,11,67] suggest that the processes that occur in lipid droplets play an important role in the molecular mechanisms of brain carcinogenesis. The localization of retinol binding protein I in lipid droplets associated with retinol accumulation suggests that lipid droplets are involved in a chain of events starting from the entry of retinol into the interior of the cell to the final stage of induction of target gene expression in the nucleus. Although the role of vitamin A in these processes remains poorly understood, many studies together with the results presented in this paper will shed light on its role in carcinogenesis.

Our results show the interplay between retinol, retinol binding proteins, cytochrome c and enhanced de novo lipid synthesis.

Considering the results presented in this paper, growing evidence suggests a far more fundamental role for retinol than previously reported. Our results suggest that retinol is essential both for signal transduction and metabolic reprogramming in cancer cells, and these processes depend on each other. We showed that these functions are governed and monitored by alterations in chemical composition of lipid droplets and mitochondria.

## Conclusions

This paper addresses the role of retinol, cellular retinoid-binding proteins, and cytochrome c in normal and cancer cells, especially the selective cellular uptake, intracellular transport, and metabolism of retinoids in normal and brain cancer cells.

We report a method for tracking the spatial distribution of retinoids, retinoid binding proteins and cytochrome c in cells using label-free hyperspectral Raman microscopy and multivariate chemometric analysis in normal human astrocytes and high-grade brain cancer cells of glioblastoma. Here we report the spectral identification of retinoids accumulation in specific cell organelles by label-free hyperspectral Raman microscopy and imaging. This method allows to monitor chemical composition of lipid droplets, mitochondria, and nuclei in single cells. Raman polarization measurements have revealed the occurrence of conformational changes that affect discrete regions of the protein molecule associated with retinol binding.

We discovered previously unknown two types of lipid droplets with distinct chemical compositions, biological functions and vibrational properties. The composition of the lipid droplets in cancer cells are dominated by triglycerides (TAG), both for medulloblastoma and glioblastoma tissues and glioblastoma cells (U-87 MG). The lipid droplets in normal astrocytes NHA are dominated by retinyl esters as well as their retinoid derivatives.

As the composition of the lipid droplets in normal, low-grade and high-grade malignant human brain cells showed remarkable differences we suggest that this is associated with different functions regulated by the relative contribution from TAGs and retinyl esters. These two types of lipid droplets are related to different functions - energy storage and signaling. We showed that their expression (observed by Raman method) and biochemical composition depends on cancer aggressiveness. The lipid droplets in cancer cells are predominantly filled with TAGs and are involved in energy storage. The lipid droplets in normal cells are filled mainly with retinyl esters /retinol and are involved in signaling, especially JAK2/STAT6 pathway signaling. An understanding of JAK2/STRA6 signaling is clinically relevant in cancer, but a detailed potential role for STRA6 awaits further investigation.

We found a higher concentration of the retinol binding proteins and a lower concentration of TAG lipids in NHA normal cells than in aggressive tumor cells of U-87 MG. The Raman intensity ratio 2848/2923 of the TAG bands at 2848 and 2923 cm^-1^ for mitochondria was 0.48 in NHA and 0.70 in U-87 MG, indicating a lower concentration of TAG lipids in NHA normal cells than in U-87 MG cancer cells.

Precise information regarding the role of retinoids, retinol binding proteins and cytochrome c is relevant both for basic cancer research and clinical applications. Understanding the metabolism of retinoids in normal and brain cancer cells is still very limited because so far there were no experimental methods available to track retinoids in vivo in living cells. The Raman-based method we present in this paper provides an excellent tool to extend our knowledge in this field.

## Materials and Methods

### Reference chemicals

All trans-retinal (no. R2500), 13-cis–retinol (no. R0271), retinol palmitate (no. 46959-U), retinol binding protein I from human urine (RBP-1, no. R9388, solution in 10 mM sodium phosphate, pH 7.4, 150 mM NaCl, and 0.05% NaN_3_), retinol (no. R7632), retinoic acid (np. R2625), an all trans-retinal (no. R2500), cytochrome c (no. C2506), β-carotene (no. C4582) were purchased from Sigma Aldrich and all-trans-retinyl oleate (no. sc-476306) from Santa Cruz Biotechnology.

### Cell culture and preparation for Raman/fluorescence spectroscopy

A normal human astrocyte (NHA) cell line (CC-2565; Lonza) and human glioblastoma (U-87 MG) cell line (HTB-14; ATCC) were used. NHA cells were grown in AGM BulletKit (Lonza CC-3186). U-87 MG cells were grown in EMEM with 2 mM glutamine, 1% non-essential amino acids (NEAA), 1 mM sodium pyruvate (NaP) and 10% foetal bovine serum (FBS). All cell lines were maintained at 37°C in a humidified atmosphere containing 5% CO_2_. Cells for Raman spectroscopy analysis were seeded on a 25-mm round CaF_2_ window placed in a 35-mm Petri dish at a density of 5×10^4^ cells per Petri dish.

Before the Raman measurements, the cells were washed with phosphate-buffered saline (PBS) to remove any unattached cells, fixed in neutral buffered 10% formalin and washed with PBS. After the Raman imaging measurements, the cells were exposed to Hoechst 33342 (25 μL at 1 μg/mL per mL of PBS) and Oil Red O (10 μL of 0.5 mM Oil Red dissolved in 60% isopropanol/dH2O per each mL of PBS) by incubation for 15 min. The cells were then washed with PBS, followed by the addition of fresh PBS for fluorescence imaging on an Alpha 300RSA WITec microscope. Mitochondria were stained by Mitotracker Red CMXROS (M7512, at a final concentration of 100 nM Mitotracker Red CMXROS for 15 mins).

### Tissue preparation for Raman spectroscopy

Raman spectra and images were analysed from patients with brain tumours. All of the procedures for human tissues were conducted under a protocol approved by the institutional Bioethical Committee at the Medical University of Lodz (RNN/323/17/KE/17/10/2017). All of the brain experiments were performed in compliance with the relevant laws and guidelines of the Bioethical Committee at the Polish Mother’s Memorial Hospital Research Institute in Lodz (no. 53/216). Written informed consent was obtained from all patients, or if subjects are under 18, from a parent and/or legal guardian.

The ex-vivo samples were obtained during resection surgery from the tumour mass. All tissue specimens were frozen and stored at -80 °C. Before the measurements, the frozen samples were cryosectioned at - 20 °C with a microtome (Microm HM 550, Thermo Fisher, Waltham, USA) into 16 µm thick slices and placed onto calcium fluoride windows (CaF_2_, 25 ×1 mm, Crystal GmbH, Berlin, Germany).

### Raman data acquisition and chemometric analysis

Raman spectroscopy measurements were performed with a confocal Raman microscope (Alpha 300RSA, WITec, Ulm, Germany) equipped with 355, 532 and 785 nm diode lasers, a 300-mm triple grating imaging spectrometer (Acton SpectraPro SP-2300; Princeton Instruments Inc., USA), a thermoelectrically cooled CCD camera (Andor, Ireland), and a 40x water dipping objective. Raman peak positions were checked using reference sample (a silicon wafer Raman peak at 520.7 cm^-1^). Raman maps were collected using a 0.3 s integration time (10 mW power at 532 nm excitation; 2 mW at 355 nm excitation ; 80 mW at 785 nm excitation) and 1 μm resolution step.

The configuration employed in the measurements is presented in Fig 12.

**Fig 12.**
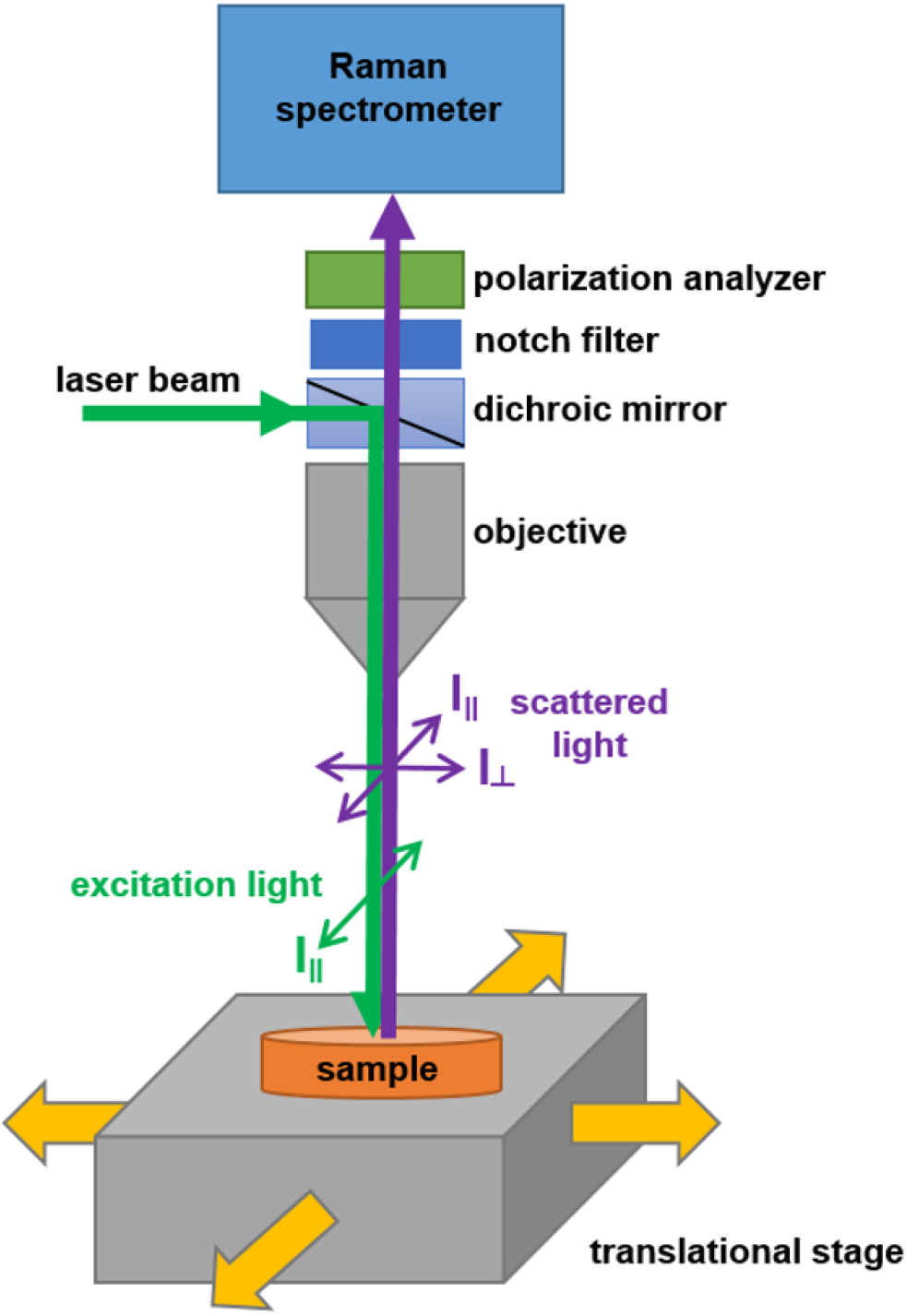
Configuration for polarized Raman imaging with an alpha 300 RSA+ (WITec, Ulm, Germany).

Following the removal of the polarization analyser of the scattered beam, we measured the Raman signal from all polarization directions (non-polarized Raman signal). When the polarization analyser was set at 0 ° with respect to the polarization of the incident excitation beam, we measured the signal of the scattered beam with polarization parallel to the polarization of the incident beam (parallel polarization III). When the polarization analyser is set at 90 ° with respect to the polarization of the incident excitation beam, we measured the signal of the scattered beam with polarization perpendicular to the polarization of the incident beam (perpendicular polarization I┴).

Data processing was performed using Project Plus Four (WITec GmbH, Ulm, Germany) and Origin 2018 (OriginLab, Northampton, USA). All Raman spectra were cosmic ray and baseline corrected (polynomial order: 3-5) and then smoothed using a Savitzky-Golay filter (order 3, 4 pt) and normalized by vector norm.

Spectroscopic data for single cells were analysed using Cluster Analysis method. Briefly Cluster Analysis is a form of exploratory data analysis in which observations are divided into different groups that have some common characteristics – vibrational features in our case. Cluster Analysis constructs groups (or classes or clusters) based on the principle that: within a group the observations must be as similar as possible, while observations belonging to different groups must be as different.

The partition of n observations (x) into k (k≤n) clusters S should be done to minimize the variance (Var) according to the formula:

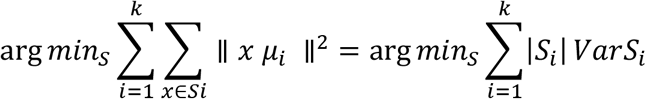

where *μ*_*i*_ is the mean of points *S*_*i*_.

K-means clustering was performed on single cells’ spectroscopic data using WITec Project Plus (WITec, Ulm, Germany). Raman images were generated by using 6 clusters (nucleus, lipid droplets, cytoplasm, mitochondria, cell border and area out of cell).

The method applied to create Raman images of tissues was the basis analysis (BBM). The basis spectra that were used in BBM represented the average spectra from characteristic morphological areas in the tissue. In BBM, each recorded spectrum of the 2D spectral Raman image was compared to the basis spectrum using a least squares fit. The Raman map illustrates the weight factor at each pixel in the 2D image coded with arbitrary colours corresponding to the selected BBM spectra. The colour code of the Raman maps was based on the integrated Raman intensities in specific regions (sum option in the filter manager in the Witec Control version 4.1 software (WITec, Ulm, Germany)).

## Author Contributions

Conceptualization: HA, JS; Funding acquisition: HA; Investigation: AI and JS; Methodology: HA, JS; Writing – original draft: HA; Writing – review & editing: HA, and JS. All authors reviewed and provide feedback on the manuscripts.

## Funding

This research was funded by Narodowe Centrum Nauki (Poland), grant number UMO-2019/33/B/ST4/01961.

## Conflicts of Interest

The authors declare no conflict of interest. The funders had no role in the design of the study; in the collection, analyses, or interpretation of data; in the writing of the manuscript, or in the decision to publish the results.

## Ethics approval and consent to participate

All of the procedures for human breast and small intestine tissues were conducted under a protocol approved by the institutional Bioethical Committee at the Medical University of Lodz (RNN/323/17/KE/17/10/2017). All of the brain experiments were performed in compliance with the relevant laws and guidelines of the Bioethical Committee at the Polish Mother’s Memorial Hospital Research Institute in Lodz (53/216). Written informed consent was obtained from all patients, or if subjects are under 18, from a parent and/or legal guardian.

## Abbreviations

apo-CRBP1: apo-cellular retinol-binding protein-I
ATP: adenosine triphosphate
CRBP-1: cellular retinol-binding protein-I
JAK: Janus kinase
LD: lipid droplets
MRI: magnetic resonance imaging
NHA: normal astrocytes (CC-2565, Lonza)
OA: oleic acid
PKC delta: protein kinase C delta
RA: retinoic acid
RBP-1: retinol binding protein I
RE: retinoids
RI: Raman imaging
ROL: retinol
RP: retinyl palmitate
STAT: signal transducer and activator of transcription
STRA6: stimulated by retinoic acid 6
TAG: trigyceride as glyceryl tripalmitate
U-87 MG: high-grade cancer cells of human glioblastoma (HTB-14, ATCC)

## Notes

### Competing Interest Statement

The authors have declared no competing interest.

## References

[1] E.F. Farias, D.E. Ong, N.B. Ghyselinck, S. Nakajo, Y.S. Kuppumbatti, R. Mira y Lopez, Cellular retinol-binding protein I, a regulator of breast epithelial retinoic acid receptor activity, cell differentiation, and tumorigenicity, J. Natl. Cancer Inst. 97 (2005) 21–29. https://doi.org/10.1093/jnci/dji004.

[2] D.C. Berry, S.M. O’Byrne, A.C. Vreeland, W.S. Blaner, N. Noy, Cross talk between signaling and vitamin A transport by the retinol-binding protein receptor STRA6, Mol. Cell. Biol. 32 (2012) 3164–3175. https://doi.org/10.1128/MCB.00505-12.

[3] J.A. Silvaroli, J.M. Arne, S. Chelstowska, P.D. Kiser, S. Banerjee, M. Golczak, Ligand Binding Induces Conformational Changes in Human Cellular Retinol-binding Protein 1 (CRBP1) Revealed by Atomic Resolution Crystal Structures, J. Biol. Chem. 291 (2016) 8528–8540. https://doi.org/10.1074/jbc.M116.714535.

[4] I. Menozzi, F. Vallese, E. Polverini, C. Folli, R. Berni, G. Zanotti, Structural and molecular determinants affecting the interaction of retinol with human CRBP1, J. Struct. Biol. 197 (2017) 330–339. https://doi.org/10.1016/j.jsb.2016.12.012.

[5] C. Folli, V. Calderone, S. Ottonello, A. Bolchi, G. Zanotti, M. Stoppini, R. Berni, Identification, retinoid binding, and x-ray analysis of a human retinol-binding protein, Proc. Natl. Acad. Sci. U. S. A. 98 (2001) 3710–3715. https://doi.org/10.1073/pnas.061455898.

[6] R.C. Moon, Comparative aspects of carotenoids and retinoids as chemopreventive agents for cancer, J. Nutr. 119 (1989) 127–134. https://doi.org/10.1093/jn/119.1.127.

[7] K.H. Dragnev, J.R. Rigas, E. Dmitrovsky, The retinoids and cancer prevention mechanisms, The Oncologist. 5 (2000) 361–368. https://doi.org/10.1634/theoncologist.5-5-361.

[8] H. Abramczyk, B. Brozek-Pluska, New look inside human breast ducts with Raman imaging. Raman candidates as diagnostic markers for breast cancer prognosis: Mammaglobin, palmitic acid and sphingomyelin, Anal. Chim. Acta. 909 (2016) 91–100. https://doi.org/10.1016/j.aca.2015.12.038.

[9] J. Surmacki, B. Brozek-Pluska, R. Kordek, H. Abramczyk, The lipid-reactive oxygen species phenotype of breast cancer. Raman spectroscopy and mapping, PCA and PLSDA for invasive ductal carcinoma and invasive lobular carcinoma. Molecular tumorigenic mechanisms beyond Warburg effect, The Analyst. 140 (2015) 2121–2133. https://doi.org/10.1039/c4an01876a.

[10] H. Abramczyk, B. Brozek-Pluska, Apical-basal polarity of epithelial cells imaged by Raman microscopy and Raman imaging: Capabilities and challenges for cancer research, J. Mol. Liq. 245 (2017) 52–61. https://doi.org/10.1016/j.molliq.2017.05.142.

[11] N. Testerink, M. Ajat, M. Houweling, J.F. Brouwers, V.V. Pully, H.-J. van Manen, C. Otto, J.B. Helms, A.B. Vaandrager, Replacement of retinyl esters by polyunsaturated triacylglycerol species in lipid droplets of hepatic stellate cells during activation, PloS One. 7 (2012) e34945. https://doi.org/10.1371/journal.pone.0034945.

[12] H. Abramczyk, B. Brozek-Pluska, A. Jarota, J. Surmacki, A. Imiela, M. Kopeć, A look into the use of Raman spectroscopy for brain and breast cancer diagnostics: linear and non-linear optics in cancer research as a gateway to tumor cell identity, Expert Rev. Mol. Diagn. https://doi.org/10.1080/14737159.2020.1724092

[13] H. Abramczyk, J. Surmacki, Antitumor Activity of Dietary Carotenoids, and Prospects for Applications in Therapy, in: Carotenoids, John Wiley & Sons, Ltd, 2016: pp. 31–42. https://doi.org/10.1002/9781118622223.ch3.

[14] D.A. Bender, Nutritional Biochemistry of the Vitamins by David A. Bender, Camb. Core. (2003). https://doi.org/10.1017/CBO9780511615191.

[15] Y.-K. Kim, L. Wassef, S. Chung, H. Jiang, A. Wyss, W.S. Blaner, L. Quadro, β-Carotene and its cleavage enzyme β-carotene-15,15′-oxygenase (CMOI) affect retinoid metabolism in developing tissues, FASEB J. 25 (2011) 1641–1652. https://doi.org/10.1096/fj.10-175448.

[16] I. of M. (US) P. on Micronutrients, Uses of Dietary Reference Intakes, National Academies Press (US), 2001. https://www.ncbi.nlm.nih.gov/books/NBK222330/

[17] M.E. Gottesman, L. Quadro, W.S. Blaner, Studies of vitamin A metabolism in mouse model systems, BioEssays News Rev. Mol. Cell. Dev. Biol. 23 (2001) 409–419. https://doi.org/10.1002/bies.1059.

[18] J.L. Napoli, Retinoic acid synthesis from beta-carotene in vitro, Methods Enzymol. 214 (1993) 193–202. https://doi.org/10.1016/0076-6879(93)14066-r.

[19] D.E. Ong, Cellular retinoid-binding proteins, Arch. Dermatol. 123 (1987) 1693–1695a.

[20] A.C. Ross, M.E. Ternus, Vitamin A as a hormone: Recent advances in understanding the actions of retinol, retinoic acid, and beta carotene, J. Am. Diet. Assoc. 93 (1993) 1285–1290. https://doi.org/10.1016/0002-8223(93)91956-Q.

[21] M. Chivot, Retinoid therapy for acne. A comparative review, Am. J. Clin. Dermatol. 6 (2005) 13–19. https://doi.org/10.2165/00128071-200506010-00002.

[22] J.M. Love, L.J. Gudas, Vitamin A, differentiation and cancer, Curr. Opin. Cell Biol. 6 (1994) 825–831. https://doi.org/10.1016/0955-0674(94)90051-5.

[23] R.M. Niles, Signaling pathways in retinoid chemoprevention and treatment of cancer, Mutat. Res. Mol. Mech. Mutagen. 555 (2004) 97–105. https://doi.org/10.1016/j.mrfmmm.2004.05.020.

[24] C.B. Stephensen, Vitamin a, Infection, and Immune Function*, Annu. Rev. Nutr. 21 (2001) 167–192. https://doi.org/10.1146/annurev.nutr.21.1.167.

[25] G.H. Travis, M. Golczak, A.R. Moise, K. Palczewski, Diseases caused by defects in the visual cycle: retinoids as potential therapeutic agents, Annu. Rev. Pharmacol. Toxicol. 47 (2007) 469–512. https://doi.org/10.1146/annurev.pharmtox.47.120505.105225.

[26] C.C. Zouboulis, Retinoids – Which Dermatological Indications Will Benefit in the Near Future?, Skin Pharmacol. Physiol. 14 (2001) 303–315. https://doi.org/10.1159/000056361.

[27] E. Doldo, G. Costanza, S. Agostinelli, C. Tarquini, A. Ferlosio, G. Arcuri, D. Passeri, M.G. Scioli, A. Orlandi, Vitamin A, cancer treatment and prevention: the new role of cellular retinol binding proteins, BioMed Res. Int. 2015 (2015) 624627. https://doi.org/10.1155/2015/624627.

[28] L. Altucci, H. Gronemeyer, The promise of retinoids to fight against cancer, Nat. Rev. Cancer. 1 (2001) 181–193. https://doi.org/10.1038/35106036.

[29] null Siddikuzzaman, C. Guruvayoorappan, V.M. Berlin Grace, All trans retinoic acid and cancer, Immunopharmacol. Immunotoxicol. 33 (2011) 241–249. https://doi.org/10.3109/08923973.2010.521507.

[30] R.M. Niles, Use of Vitamins A and D in Chemoprevention and Therapy of Cancer: Control of Nuclear Receptor Expression and Function, in: Diet Cancer Mol. Mech. Interact., Springer US, Boston, MA, 1995: pp. 1–15. https://doi.org/10.1007/978-1-4899-0949-7_1.

[31] O. Arrieta, C.H. González-De la Rosa, E. Aréchaga-Ocampo, G. Villanueva-Rodríguez, T.L. Cerón-Lizárraga, L. Martínez-Barrera, M.E.1 Vázquez-Manríquez, M.A. Ríos-Trejo, M.A. Alvarez-Avitia, N. Hernández-Pedro, C. Rojas-Marín, J. De la Garza, Randomized phase II trial of All-trans-retinoic acid with chemotherapy based on paclitaxel and cisplatin as first-line treatment in patients with advanced non-small-cell lung cancer, J. Clin. Oncol. Off. J. Am. Soc. Clin. Oncol. 28 (2010) 3463–3471. https://doi.org/10.1200/JCO.2009.26.6452.

[32] M. Bryan, E.D. Pulte, K.C. Toomey, L. Pliner, A.C. Pavlick, T. Saunders, R. Wieder, A pilot phase II trial of all-trans retinoic acid (Vesanoid) and paclitaxel (Taxol) in patients with recurrent or metastatic breast cancer, Invest. New Drugs. 29 (2011) 1482–1487. https://doi.org/10.1007/s10637-010-9478-3.

[33] A.R. Mawson, Retinoids in the treatment of glioma: a new perspective, Cancer Manag. Res. 4 (2012) 233–241. https://doi.org/10.2147/CMAR.S32449.

[34] J.L. Napoli, Cellular retinoid binding-proteins, CRBP, CRABP, FABP5: Effects on retinoid metabolism, function and related diseases, Pharmacol. Ther. 173 (2017) 19–33. https://doi.org/10.1016/j.pharmthera.2017.01.004.

[35] R. Acin-Perez, B. Hoyos, F. Zhao, V. Vinogradov, D.A. Fischman, R.A. Harris, M. Leitges, N. Wongsiriroj, W.S. Blaner, G. Manfredi, U. Hammerling, Control of oxidative phosphorylation by vitamin A illuminates a fundamental role in mitochondrial energy homoeostasis, FASEB J. Off. Publ. Fed. Am. Soc. Exp. Biol. 24 (2010) 627–636. https://doi.org/10.1096/fj.09-142281.

[36] O. Warburg, On the Origin of Cancer Cells, Science. 123 (1956) 309–314. https://doi.org/10.1126/science.123.3191.309.

[37] A. Schulze, J. Downward, Flicking the Warburg switch-tyrosine phosphorylation of pyruvate dehydrogenase kinase regulates mitochondrial activity in cancer cells, Mol. Cell. 44 (2011) 846–848. https://doi.org/10.1016/j.molcel.2011.12.004.

[38] T. Hitosugi, S. Kang, M.G. Vander Heiden, T.-W. Chung, S. Elf, K. Lythgoe, S. Dong, S. Lonial, X. Wang, G.Z. Chen, J. Xie, T.-L. Gu, R.D. Polakiewicz, J.L. Roesel, T.J. Boggon, F.R. Khuri, D.G. Gilliland, L.C. Cantley, J. Kaufman, J. Chen, Tyrosine phosphorylation inhibits PKM2 to promote the Warburg effect and tumor growth, Sci. Signal. 2 (2009) ra73. https://doi.org/10.1126/scisignal.2000431.

[39] H. Abramczyk, A. Imiela, B. Brożek-Płuska, M. Kopeć, J. Surmacki, A. Śliwińska, Aberrant Protein Phosphorylation in Cancer by Using Raman Biomarkers, Cancers. 11 (2019) 2017. https://doi.org/10.3390/cancers11122017.

[40] J. Kwong, K.-W. Lo, L.S.-N. Chow, K.-F. To, K.-W. Choy, F.L. Chan, S.C. Mok, D.P. Huang, Epigenetic silencing of cellular retinol-binding proteins in nasopharyngeal carcinoma, Neoplasia N. Y. N. 7 (2005) 67–74. https://doi.org/10.1593/neo.04370.

[41] N. Noy, Vitamin A Transport and Cell Signaling by the Retinol-Binding Protein Receptor STRA6, Subcell. Biochem. 81 (2016) 77–93. https://doi.org/10.1007/978-94-024-0945-1_3.

[42] W.S. Blaner, STRA6, a cell-surface receptor for retinol-binding protein: the plot thickens, Cell Metab. 5 (2007) 164–166. https://doi.org/10.1016/j.cmet.2007.02.006.

[43] R. Goswami, M.H. Kaplan, STAT Transcription Factors in T Cell Control of Health and Disease, Int. Rev. Cell Mol. Biol. 331 (2017) 123–180. https://doi.org/10.1016/bs.ircmb.2016.09.012.

[44] A. Imiela, B. Polis, L. Polis, H. Abramczyk, Novel strategies of Raman imaging for brain tumor research, Oncotarget. 8 (2017) 85290–85310. https://doi.org/10.18632/oncotarget.19668.

[45] H. Abramczyk, J. Surmacki, M. Kopeć, A.K. Olejnik, K. Lubecka-Pietruszewska, K. Fabianowska-Majewska, The role of lipid droplets and adipocytes in cancer. Raman imaging of cell cultures: MCF10A, MCF7, and MDA-MB-231 compared to adipocytes in cancerous human breast tissue, The Analyst. 140 (2015) 2224–2235. https://doi.org/10.1039/c4an01875c.

[46] H. Abramczyk, J. Surmacki, M. Kopeć, A.K. Olejnik, A. Kaufman-Szymczyk, K. Fabianowska-Majewska, Epigenetic changes in cancer by Raman imaging, fluorescence imaging, AFM and scanning near-field optical microscopy (SNOM). Acetylation in normal and human cancer breast cells MCF10A, MCF7 and MDA-MB-231, The Analyst. 141 (2016) 5646–5658. https://doi.org/10.1039/c6an00859c.

[47] C.-S. Liao, M.N. Slipchenko, P. Wang, J. Li, S.-Y. Lee, R.A. Oglesbee, J.-X. Cheng, Microsecond Scale Vibrational Spectroscopic Imaging by Multiplex Stimulated Raman Scattering Microscopy, Light Sci. Appl. 4 (2015). https://doi.org/10.1038/lsa.2015.38.

[48] H. Abramczyk, A. Imiela, The biochemical, nanomechanical and chemometric signatures of brain cancer, Spectrochim. Acta. A. Mol. Biomol. Spectrosc. 188 (2018) 8–19. https://doi.org/10.1016/j.saa.2017.06.037.

[49] M. Ji, D.A. Orringer, C.W. Freudiger, S. Ramkissoon, X. Liu, D. Lau, A.J. Golby, I. Norton, M. Hayashi, N.Y.R. Agar, G.S. Young, C. Spino, S. Santagata, S. Camelo-Piragua, K.L. Ligon, O. Sagher, X.S. Xie, Rapid, label-free detection of brain tumors with stimulated Raman scattering microscopy, Sci. Transl. Med. 5 (2013) 201ra119. https://doi.org/10.1126/scitranslmed.3005954.

[50] S. Berezhna, H. Wohlrab, P.M. Champion, Resonance Raman investigations of cytochrome c conformational change upon interaction with the membranes of intact and CA2+-exposed mitochondria, Biochemistry. 42 (2003) 6149–6158. https://doi.org/10.1021/bi027387y.

[51] F. Adar, M. Erecinska, University of Pennsylvania, Resonance Raman spectra of whole mitochondria, Biochemistry. 17 (1978) 5484–5488. https://doi.org/10.1021/bi00618a024.

[52] M. Okada, N.I. Smith, A.F. Palonpon, H. Endo, S. Kawata, M. Sodeoka, K. Fujita, Label-free Raman observation of cytochrome c dynamics during apoptosis, Proc. Natl. Acad. Sci. 109 (2012) 28–32. https://doi.org/10.1073/pnas.1107524108.

[53] C. Onogi, H. Hamaguchi, In vivo Detection of Ferrous Cytochrome c in Mitochondria of Single Living Yeast Cells by Resonance Raman microspectroscopy, AIP Conf. Proc. 1267 (2010) 362. https://doi.org/10.1063/1.3482559.

[54] J.M. Surmacki, B.J. Woodhams, A. Haslehurst, B.A.J. Ponder, S.E. Bohndiek, Raman micro-spectroscopy for accurate identification of primary human bronchial epithelial cells, Sci. Rep. 8 (2018) 1–11. https://doi.org/10.1038/s41598-018-30407-8.

[55] Reference database of Raman spectra of biological molecules - De Gelder - 2007 - Journal of Raman Spectroscopy - Wiley Online Library, (n.d.). https://onlinelibrary.wiley.com/doi/abs/10.1002/jrs.1734 (accessed October 30, 2019).

[56] J.M. Surmacki, Monitoring the effect of therapeutic doses of gamma irradiation on medulloblastoma by Raman spectroscopy, Anal. Methods. 12 (2020) 383–391. https://doi.org/10.1039/C9AY02238D.

[57] H. Abramczyk, A. Imiela, B. Brozek-Pluska, M. Kopec, Advances in Raman imaging combined with AFM and fluorescence microscopy are beneficial for oncology and cancer research, Nanomed. 14 (2019) 1873–1888. https://doi.org/10.2217/nnm-2018-0335.

[58] H. Abramczyk, J.M. Surmacki, B. Brozek-Pluska, Redox state changes of mitochondrial cytochromes in brain and breast cancers by Raman spectroscopy and imaging, BioRxiv. (2020) 2020.05.08.083915. https://doi.org/10.1101/2020.05.08.083915.

[59] P.T. Bozza, J.P.B. Viola, Lipid droplets in inflammation and cancer, Prostaglandins Leukot. Essent. Fatty Acids. 82 (2010) 243–250. https://doi.org/10.1016/j.plefa.2010.02.005.

[60] P.P. Fu, Q. Xia, J.J. Yin, S.-H. Cherng, J. Yan, N. Mei, T. Chen, M.D. Boudreau, P.C. Howard, W.G. Wamer, Photodecomposition of Vitamin A and Photobiological Implications for the Skin†, Photochem. Photobiol. 83 (2007) 409–424. https://doi.org/10.1562/2006-10-23-IR-1065.

[61] N. Stone, C. Kendall, J. Smith, P. Crow, H. Barr, Raman spectroscopy for identification of epithelial cancers, Faraday Discuss. 126 (2004) 141–157; discussion 169-183. https://doi.org/10.1039/b304992b.

[62] G.J. Thomas, Raman spectroscopy of protein and nucleic acid assemblies, Annu. Rev. Biophys. Biomol. Struct. 28 (1999) 1–27. https://doi.org/10.1146/annurev.biophys.28.1.1.

[63] J.M. Shaver, K.A. Christensen, J.A. Pezzuti, M.D. Morris, Structure of Dihydrogen Phosphate Ion Aggregates by Raman-Monitored Serial Dilution, Appl. Spectrosc. 52 (1998) 259–264.

[64] H. Deng, V.A. Bloomfield, J.M. Benevides, G.J. Thomas, Dependence of the Raman signature of genomic B-DNA on nucleotide base sequence, Biopolymers. 50 (1999) 656–666. https://doi.org/10.1002/(SICI)1097-0282(199911)50:6<656::AID-BIP10>3.0.CO;2-9.

[65] Y. Guan, G.S.-C. Choy, R. Glaser, G.J. Thomas, Vibrational Analysis of Nucleic Acids. 2. Ab Initio Calculation of the Molecular Force Field and Normal Modes of Dimethyl Phosphate, J. Phys. Chem. 99 (1995) 12054–12062. https://doi.org/10.1021/j100031a039.

[66] H. Abramczyk, Femtosecond primary events in bacteriorhodopsin and its retinal modified analogs: revision of commonly accepted interpretation of electronic spectra of transient intermediates in the bacteriorhodopsin photocycle, J. Chem. Phys. 120 (2004) 11120–11132. https://doi.org/10.1063/1.1737731.

[67] I.Y. Benador, M. Veliova, K. Mahdaviani, A. Petcherski, J.D. Wikstrom, E. Assali, R. Acín-Peréz, M. Shum, M.F. Oliveira, S. Cinti, C. Sztalryd, W.D. Barshop, J.A. Wohlschlegel, B.E. Corkey, M. Liesa, O.S. Shirihai, Mitochondria Bound to Lipid Droplets Have Unique Bioenergetics, Composition, and Dynamics That Support Lipid Droplet Expansion, Cell Metab. 27 (2018) 869-885.e6. https://doi.org/10.1016/j.cmet.2018.03.003.

[68] M.E. Newcomer, T.A. Jones, J. Aqvist, J. Sundelin, U. Eriksson, L. Rask, P.A. Peterson, The three-dimensional structure of retinol-binding protein, EMBO J. 3 (1984) 1451–1454.

